# NFYA regulates two genome-wide transcriptional activation events during oocyte-to-embryo transition

**DOI:** 10.64898/2026.03.30.715371

**Authors:** Qianying Yang, Shan Jiang, Boyan Wang, Yi Zhang

## Abstract

Primordial follicle oocyte activation (PFA) and zygotic genome activation (ZGA) represent two major waves of transcription activation respectively required for oocyte growth and embryo development. Although many shared molecular hallmarks between PFA and ZGA suggest potential common mechanisms driving both waves of transcriptional activation, such regulators are yet to be identified. Here we demonstrate that the pioneer factor NFYA belongs to such regulators. Oocyte-specific *Nfya-*KO impairs open chromatin establishment and transcriptional activation during PFA, which triggers ferroptosis-like cell death leading to folliculogenesis failure. Moreover, acute NFYA-depletion downregulates 38.5% of major ZGA genes and arrests 50% of embryos at the two-three cell stage. Mechanistically, although NFYA exhibits distinct chromatin-binding preferences predominantly targeting promoters during PFA and enhancers during ZGA, pre-occupied NFYA regulates chaperone and histone genes in both events through promoter binding. Chaperone inhibition impairs PFA and ZGA. Together, we establish NFYA as a common regulator of PFA and ZGA.

## Introduction

Neonatal mammalian non-growing oocytes are arrested in the diplotene stage of meiotic prophase I and maintained in a dormant state in primordial follicles ^1^, characterized by widespread low transcriptional activity relative to growing oocytes ^2,3,5^. During primordial follicle oocyte activation (PFA), transcriptional activity increases sequentially to support oocyte growth and progression to primary follicle stage and then secondary follicle stage, termed the primordial-to-primary transition (PPT) and primary-to-secondary transition (PST), respectively ^7,8^. Oocytes thereby accumulate maternal RNAs and proteins required for subsequent oocyte growth and early embryonic development ^7,8^. Studies on the mechanisms of quiescent primordial follicle maintenance have led to the identification of several regulators, including PTEN-PI3K ^9^, FOXO3A ^10^, and LHX8 ^12,14^. Knockout of these regulators causes excessive activation and premature exhaustion of the primordial follicle pool. However, the pioneer factors responsible for initiating PFA *in vivo* have remained elusive.

Secondary follicle oocytes continue to grow and accumulate maternal factors to develop into full-grown oocytes, accompanied by a decline in transcription to quiescence, and arrest at MII stage ^7,8^. Following fertilization, the silenced genome reactivates through zygotic genome activation (ZGA), enabling the maternal-to-zygotic transition (MZT) for early embryogenesis ^16^. ZGA is initiated at the late-zygotic stage with low levels of transcription and peaks at the late 2-cell (L2C) stage with large-scale transcription ^18^. Notably, PFA and ZGA are both characterized by ribosome biogenesis, rapid and massive protein synthesis and proper folding, and widespread chromatin remodeling ^20,22,24,26^. Although these shared features are recognized, a common pioneer factor triggering both PFA and ZGA remains to be identified, largely owing to sample and technical limitations.

In this study, we demonstrate that the pioneer factor NFYA is a common regulator for both PFA and ZGA by utilizing an oocyte-specific knockout mouse model ^28^ and a targeted protein degradation tag (dTAG) mouse model ^30,31^, combined with RNA-seq, ATAC-seq, and low-input CUT&RUN assays. Phenotypically, oocyte-specific *Nfya* deletion impairs open chromatin establishment and transcriptional activation during PFA, triggering ferroptosis-like cell death and early folliculogenesis failure. By generating NFYA^dTAG^ mice, we further demonstrate that acute NFYA depletion in zygotes downregulates 38.5% of the major ZGA genes and arrests 50% of embryos at the 2-3 cell stage. Mechanistically, although NFYA exhibits distinct chromatin-binding preferences predominantly targeting promoters during PFA and enhancers during ZGA, pre-occupied NFYA regulates chaperones and histone genes in both PFA and ZGA through conserved promoter binding. Inhibition of chaperones impairs both PFA and ZGA supporting their requirement for transcriptional activation. Together, our study advances the understanding of how the same transcription factor mediates genome-wide transcriptional activation in two critical developmental transitions.

## Results

### Identification of NFYA as a potential common regulator for PFA and MZT

To identify potential regulators of oocyte-to-embryo transition, we integrated our total RNA-seq and ATAC–seq datasets with previously published data, covering developmental stages from primordial follicle oocytes to 2-cell embryos ^22,33,35,37^. Principal components analysis (PCA) revealed that both chromatin accessibility and transcription undergo two major transitions at PFA and MZT (**Figure 1A and S1A)**. Notably, chromatin remodeling in PFA was as extensive as in MZT, during which ZGA occurs ^18^. Additionally, the lower global chromatin accessibility in primordial follicle oocytes was in line with its transcriptional quiescent state compared to that of growing-oocytes **(Figure S1B-C)**. To identify candidate transcription factors (TFs) participating in both PFA and MZT, we analyzed the chromatin accessibility changes during these two transitions. We identified 109,129 newly accessible chromatin regions (P2) during primordial-to-primary transition (**Figure 1B**), while similar analysis identified 3,806 and 5,084 newly accessible chromatin regions at zygote stage (Z2 and Z3) and late 2-cell (L2C) stage (Z4), respectively (**Figure 1C**). Motif enrichment analysis of these regions revealed that OBOX ^39^, NR5A2 ^41,43^, and GABPA ^31^ are exclusively enriched during MZT, while LHX ^14^ and TCF3/12 ^44^ motifs are specifically enriched during PFA, generally aligning with their expression dynamics (**Figure 1D and S1D-F**). Notably, NFYA binding motifs are highly enriched (p < 1 × 10 ^100^) in the newly accessible regions during both PFA and MZT (NFYA motif ranked #22 in P2, #7 in Z2, and #1 in Z3), suggesting that NFYA may act as a common regulator orchestrating these two transitions.

**Figure 1.**
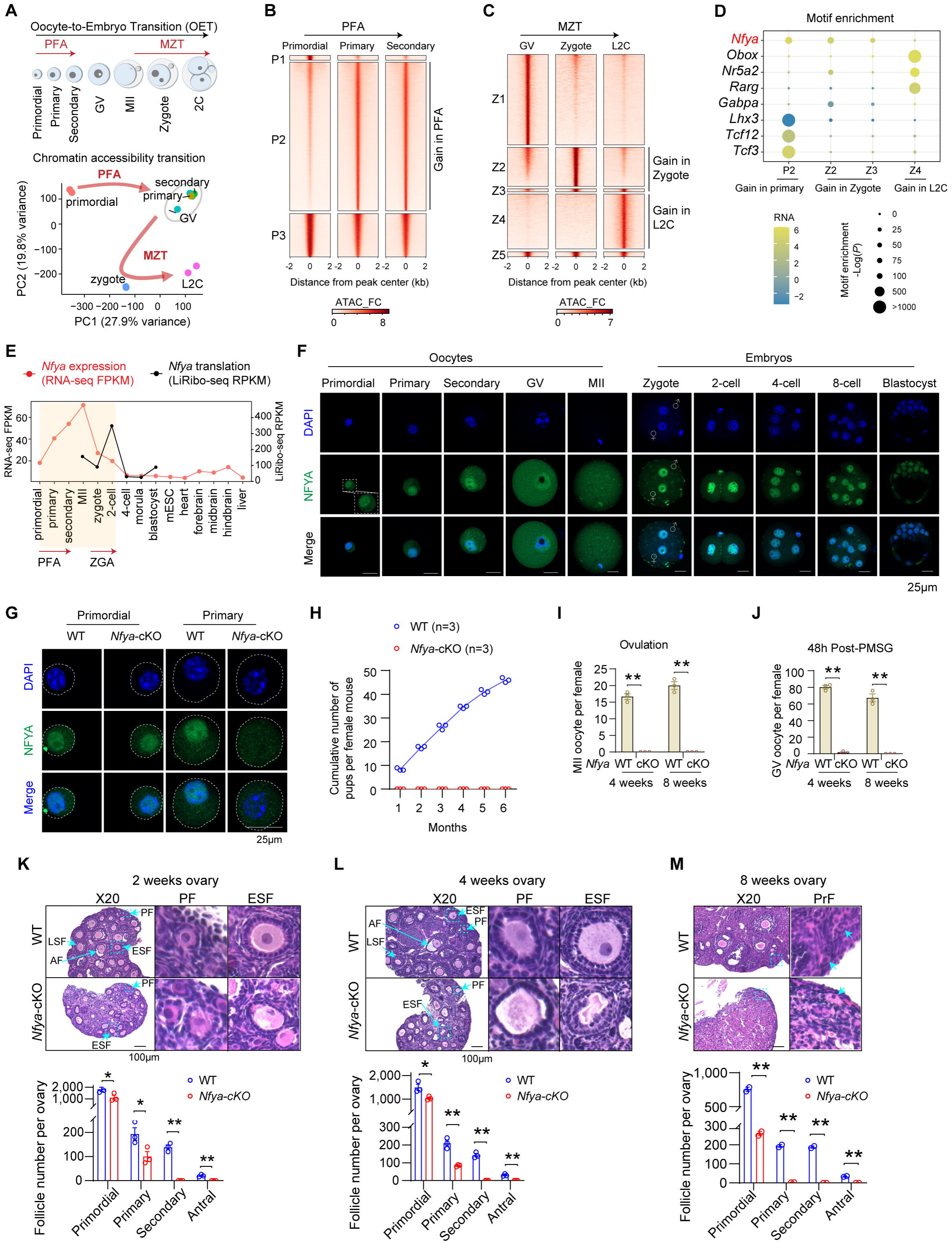
Identification of NFYA as a potential regulator of both PFA and ZGA and loss of NFYA causes folliculogenesis failure. **(A)** Upper panel: Schematic diagram of the oocyte-to-embryo transition, from the primordial follicle stage to the 2-cell embryo (2C) stage, with PFA and ZGA timing indicated. GV, germinal vesicle oocyte. Lower panel: principal component analysis (PCA) of global chromatin accessibility across different stages from primordial follicle to late 2-cell. **(B)** Heatmaps showing the ATAC-seq peaks classified based on their changes during PFA (P1-P3). P1 cluster (n=1,789) represents specific open chromatin in primordial follicle stage; P2 cluster (n=109,129) represents those that gained open chromatin in primary and secondary follicle stages; P3 cluster (n=21,717) indicates the shared open chromatin across PFA. **(C)** Heatmaps showing the ATAC-seq peaks classified based on their changes during MZT (Z1-Z5). Z1 cluster (n=8,467) represents specific open chromatin in GV; Z2 cluster (n=3,650) represents the gained open chromatin in zygote; Z3 cluster (n=156) represents the gained open chromatin in both zygote and late 2-cell (L2C); Z4 cluster (n=5,048) represents the gained open chromatin in L2C; Z5 cluster (n=249) represents the shared open chromatin across MZT. **(D)** Enrichment of representative TF motifs (from figure S1D) at the ATAC–seq peak groups (P2, Z2, Z3, and Z4) during PFA and MZT. **(E)** The dynamics of *Nfya* expression at transcription (RNA-seq, FPKM) and translation (LiRibo-seq ^22^, RPKM) levels in mouse oocytes, early embryos, and tissues. **(F)** Immunostaining of NFYA (green) during mouse oocyte and early embryo development. Scale bar, 25 μm. Dashed white box indicates a higher-magnification view of the primordial follicle oocyte. **(G)** Immunostaining of NFYA (green) in WT and *Nfya*-cKO primordial/primary follicle oocytes. Scale bar, 25 μm. The white dashed circle marks the whole oocytes. **(H)** Fertility test showing the cumulative numbers of pups from crosses of WT (n=3) and *Nfya*-cKO (n=3) females with WT males. **(I)** Numbers of MII oocytes per female mice at 4 or 8 weeks-age after superovulation (WT, n=3; *Nfya*-cKO, n=3). **(J)** Numbers of GV oocytes per female mice at 4 or 8 weeks-age after 48h PMSG induction (WT, n=3; *Nfya*-cKO, n=3). (**K-M),** Upper panel: H&E staining of paraffin embedded ovarian sections showing the histology of WT and *Nfya*-cKO ovaries at the indicated postnatal days. Scale bar, 100 μm. A high-resolution view of the boxed area is shown in parallel. Arrows point to follicles at the indicated stages. PrF, primordial follicle; PF, primary follicle; ESF, early secondary follicle; LSF, late secondary follicle; AF, antral follicle. Lower panel: Bar chart showing the follicle numbers of WT (blue) and *Nfya*-cKO (red) ovaries at the indicated times. Quantitative data are shown as mean ± SEM; ∗∗p < 0.01, ∗p < 0.05, ns p > 0.05, Student’s t test.

Consistent with this notion, NFYA transcription increases dramatically during PFA, peaks at the MII oocyte stage, and declines after fertilization, whereas its translation remains high levels throughout MZT and increases during ZGA (**Figure 1E**). Notably, *Nfya* is highly expressed during PFA and ZGA, but at a very low level in later embryos, mESCs, and somatic tissues. Immunostaining showed that NFYA protein level increases during PFA and stays in nucleus from primordial to GV stages, and translocated from the cytoplasm into the nucleus during MZT (**Figure 1F**). These data are consistent with a potential role of NFYA in regulating both PFA and MZT. Our previous studies suggested that NFYA regulates ZGA and preimplantation development ^24^. Therefore, we first focused on NFYA’s role in PFA.

### Loss of NFYA causes early folliculogenesis failure

NFYA is the sequence-specific DNA binding subunit of the NFY/CCAAT complex with an evolutionarily conserved function in chromatin accessibility establishment or maintenance ^45,46^. Global *Nfya* knockout is embryonic lethal ^47^. To study its potential role in PFA, we generated *Gdf9-Cre Nfya*^flox/flox^ mice (referred to as *Nfya*-cKO hereafter) which depletes NFYA just as primordial follicle oocyte activation begins ^28^ (**Figure S1G**). NFYA was depleted in primary but not dormant primordial follicle oocytes (**Figure 1G and S1H**). *Nfya*-cKO mice are infertile (**Figure 1H**) and cannot generate mature or full-grown oocytes (**Figure 1I-J**), suggesting early folliculogenesis defects. Furthermore, significant ovarian developmental defects were observed in *Nfya*-cKO mice starting from 2-week (**Figure S1I-J**), corresponding to the stage when the first wave of antral follicle formation is completed in WT ovaries ^7,8^. *Nfya* ablation caused a drastic loss of primary follicles (PF); some progressed to early secondary follicle (ESF) with abnormal morphology but failed to generate late secondary follicles (LSF) or antral follicles (AF) (**Figure 1K**). Notably, folliculogenesis defects are more severe in 4-week *Nfya*-cKO mice, with shrunken morphology of primary and early secondary follicle oocytes and significant loss of primary follicles, suggesting oocyte arrest and follicle degeneration and resorption (**Figure 1L**). By 8 weeks, almost all the primary follicles were degenerated in *Nfya*-cKO mice, with only primordial follicles (PrF) remaining, which are characterized by flat squamous pre-granulosa cells (**Figure 1M and S1K**), indicating defective PFA in *Nfya*-cKO mice. Taken together, these results demonstrate that NFYA is critical for early folliculogenesis.

### NFYA deficiency impairs PFA in primary follicle oocytes

To assess PFA, we performed RNA-seq for *Nfya*-cKO and WT oocytes at primordial, primary, and secondary follicle stages (**Figure S2A**). Oocyte marker genes were highly expressed and granulosa cell markers were undetectable (**Figure S2B**), indicating no granulosa cell contamination. Consistent with protein depletion (**Figure 1G**), *Nfya* RNA was depleted at primary and secondary, but not dormant primordial, follicle stages (**Figure S2C**). PCA showed that while primordial follicle oocytes of WT and *Nfya*-cKO were clustered closely, the primary and secondary follicle oocytes of *Nfya*-cKO and WT were distinctively clustered (**Figure 2A**), suggesting extensive transcriptomic alterations in the *Nfya*-cKO primary and secondary follicle oocytes. Comparative analysis identified 809 downregulated and 281 upregulated genes in primary follicle oocytes in response to NFYA depletion (**Figure 2B and Table S1**). To identify differentially expressed genes (DEGs) directly targeted by NFYA, we performed CUT&RUN experiments using WT primary follicle oocytes and confirmed that NFYA peaks are highly enriched for NFYA binding motif (**Figure S2D-E**). Importantly, NFYA CUT&RUN in *Nfya*-cKO primary follicle oocytes revealed that all the NFYA peaks were diminished compared to that in WT control (**Figure S2F**), confirming the specificity of the CUT&RUN signal. Integrative analysis showed that NFYA mainly occupies promoters of the downregulated genes with 402 out of the 809 genes bound by NFYA (**Figure 2C and Table S1**). Consistently, 95.2% (383/402) of the NFYA bound promoters contain at least one NFYA motif (**Figure 2C and S2G**). Gene Ontology (GO) analysis revealed that these direct targets are enriched for cell cycle transition and chromosome condensation (**Figure 2D**). Notably, the direct targets include many genes whose knockout or mutation causes infertility, such as *Fancf*, *Hsp90b1*, *Rfwd3*, and *Prkaca* (**Figure 2E and S2H**). Interestingly, *Prkaca* encodes the catalytic subunit of cAMP-dependent protein kinase A (PKA), and the cAMP-PKA signaling is pivotal for PFA and meiotic cell cycle arrest ^48,49^, supporting a critical role of NFYA in PFA.

**Figure 2.**
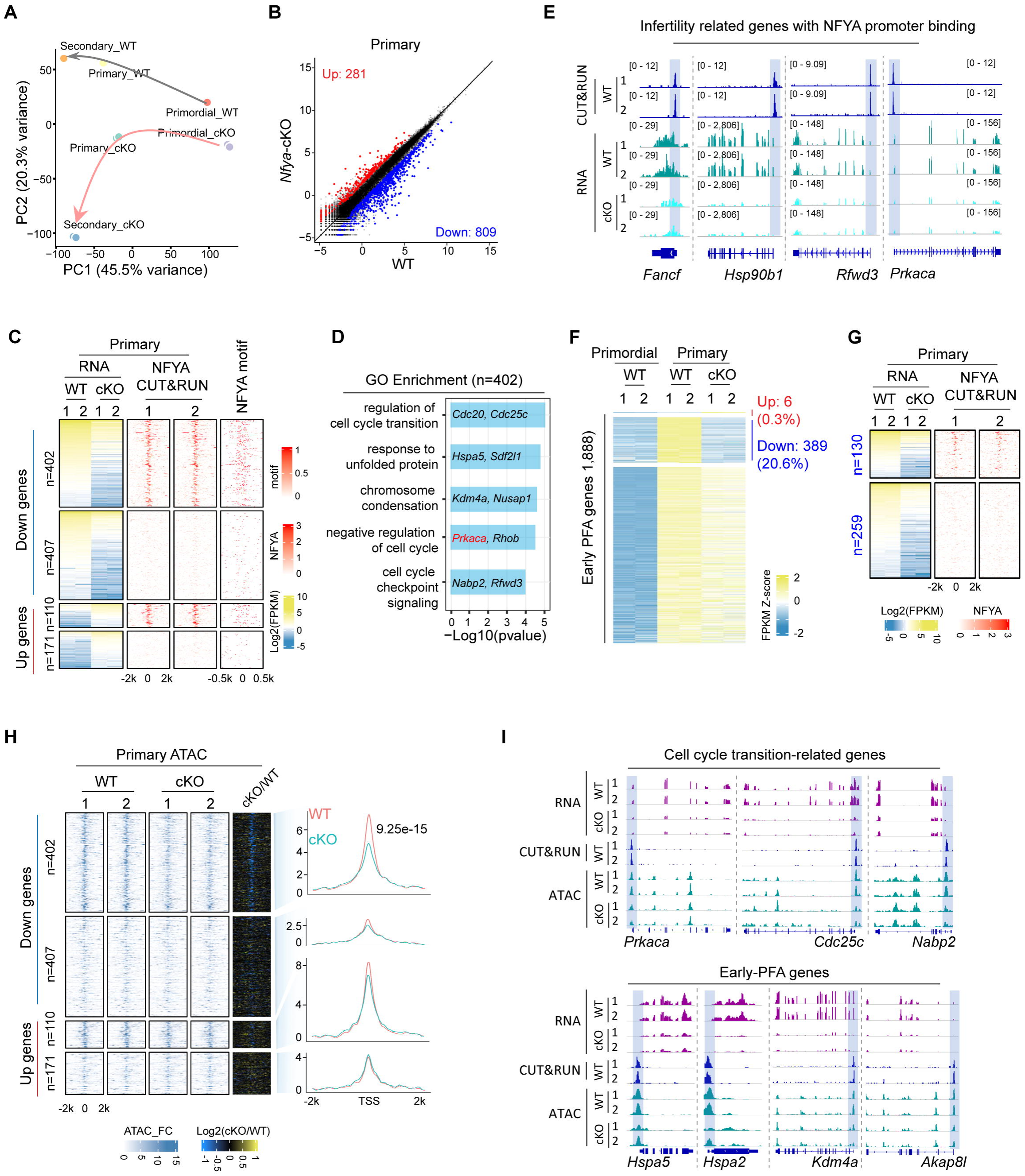
NFYA deficiency impairs PFA in primary follicle oocytes. **(A)** PCA of WT and Nfya-cKO primary and secondary follicle oocytes. **(B)** Scatter plots comparing the gene expression profiles of WT and *Nfya*-cKO primary follicle oocytes from P7 ovaries. The x and y axis of the dot plots are Log_2_CPM (counts per million) from RNA-seq. Fold change > 2, false discovery rate (FDR) < 0.05. **(C)** Heatmaps showing the DEGs of primary follicles caused by NFYA loss, as well as NFYA binding and NFYA motif occurrence profiles around the TSS of the corresponding genes. The downregulated and upregulated genes are each separated into two groups based on whether they have direct NFYA binding. **(D)** Gene Ontology (GO) terms enriched for the NFYA-bound down-regulated genes (n=402) with examples. **(E)** Genome browser view of representative NFYA-targeted infertility-related genes showing NFYA CUT&RUN and RNA-seq in WT and CKO primary follicle oocytes. **(F)** Heatmap showing early-PFA gene expression (z-scores) in WT and *Nfya*-cKO oocytes at the indicated stages. **(G)** Heatmaps showing the RNA-seq results of the downregulated early-PFA genes (n=389) at primary follicle stage upon NFYA loss, as well as the NFYA binding and NFYA motif occurrence around the TSS of the corresponding genes. The downregulated genes are separated into two groups based on whether they have direct NFYA binding. **(H)** Heatmaps and density plots showing the enrichment and average ATAC–seq signal fold change (ATAC_FC) in the promoter regions of DEGs of primary follicles upon NFYA loss, respectively. The downregulated or upregulated genes are separated into two groups based on whether they have direct NFYA promoter binding. P values are calculated with two-sided Student’s t-test. **(I)** Examples of genome browser view of NFYA binding (shaded), ATAC and RNA-seq signals in WT and cKO primary follicles.

To understand how NFYA regulates PFA, we compared the transcriptomes of primordial and primary follicle oocytes and identified 1,888 genes activated during the primordial-to-primary transition (PPT), termed early-PFA genes (**Figure 2F and Table S2**). Of these early-PFA genes, 389 (20.6%) failed to be activated in response to NFYA depletion (**Figure 2F**), accounting for 48.1% of the 809 downregulated genes (**Figure S2I**). A global disruption of early-PFA genes expression in *Nfya*-cKO primary follicle oocytes was further demonstrated by gene set enrichment analysis (GSEA) (**Figure S2J**). Among the 389 affected early-PFA genes, 130 gene promoters are directly bound by NFYA (**Figure 2G**), including *Ezh2*, *Kdm4a*, and *Sdl2l1* (**Figure S2K**). Since low-input CUT&RUN data can inherently miss peaks leading to underestimation, we performed motif analysis on the remaining 259 genes and found that 79.9% (207/259) contained at least one NFYA motif in their promoter regions (**Figure S2L**). Collectively, these data suggest a direct role of NFYA in PFA.

As NFYA is a pioneer factor essential for establishing open chromatin ^24,45^, we next asked whether NFYA regulates early-PFA genes by promoting their chromatin accessibility. Comparative ATAC-seq analysis (**Figure S2M**) revealed that NFYA loss significantly reduced chromatin accessibility at the promoters of NFYA-bound downregulated genes (for example *Prkaca*, *Hspa2*, *Kdm4a*, and *Akap8l*) (**Figure 2H-I**). Thus, NFYA contributes to promoter chromatin accessibility, and its loss causes defective PFA during primordial-to-primary follicle transition.

### NFYA deficiency impairs PFA in secondary follicle oocytes

After reaching primary follicle, PFA continues into secondary follicles to rapidly accumulate maternal mRNAs and proteins required for subsequent oocyte growth and development ^2,3^. We asked whether NFYA loss would affect gene activation in secondary follicle oocytes. To this end, transcriptomic analysis was performed, which revealed 1,844 downregulated and 1,517 upregulated genes in secondary follicle oocytes in response to NFYA loss (**Figure 3A and Table S3**). To identify the direct effects caused by NFYA loss, we performed NFYA CUT&RUN in secondary WT follicle oocytes (**Figure S3A**) and found that NFYA mainly occupies the promoter regions with highly enriched for NFYA binding motif (**Figure S3B-C**). We found that 1,064 out of the 1,844 down-regulated genes bound by NFYA (**Figure S3D and Table S3**), which is consistent with its motif enrichment in these promoters (**Figure S3D**). Since loss function of NFYA affects PFA in primary follicle oocytes (**Figure 2F**), to identify the genes specifically affected in the secondary follicle oocytes, we separated the affected genes into two groups: those affected in both primary and secondary follicle oocytes, and those affected only in secondary follicle oocytes (**Figure S3E**). Of the 1,231 secondary-specific downregulated genes, 658 are bound by NFYA with 87.5% (576/658) affected genes contain at least one NFYA binding motif in their promoters (**Figure 3B and S3F**). GO analysis revealed that these downregulated direct targets are enriched for meiotic cell cycle, response to endoplasmic reticulum (ER) stress, and chromosome segregation (**Figure S3G**). Notably, these secondary-specific direct targets include many genes whose knockout or mutation exhibit infertility phenotype such as *Calr*, *Swsap1*, *Nlrp14*, and *Piwil2* (**Figure 3C and S3H**). For example, CALR is important for quality control of nascent proteins in the ER and for oocyte development ^50^ and *Piwil2* are important for functional oocyte generation ^51,52^, consistent with a critical role of NFYA in oocyte development.

**Figure 3.**
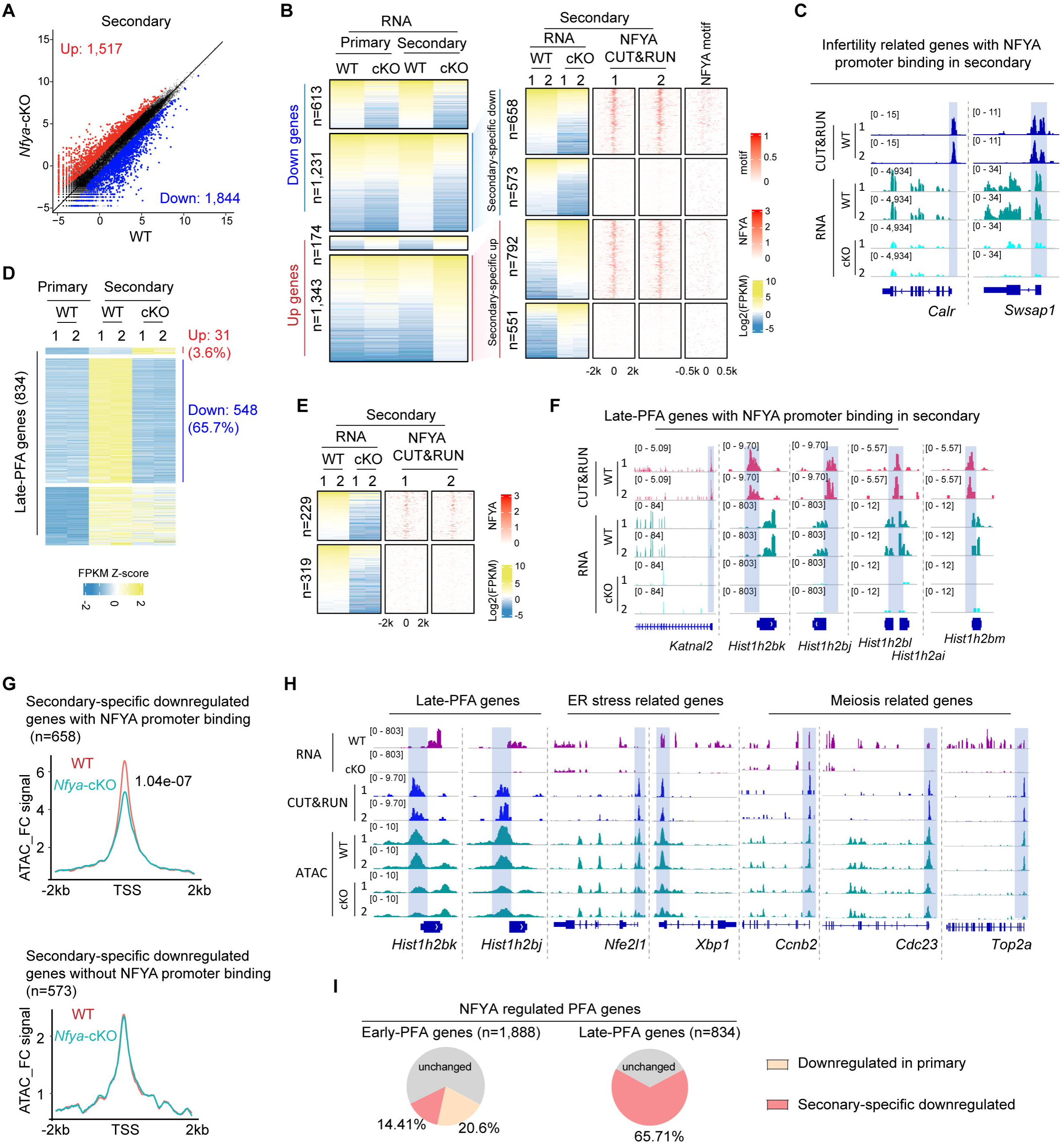
NFYA deficiency impairs PFA in secondary follicle oocytes. **(A)** Scatter plots comparing the gene expression profiles of WT and *Nfya*-cKO secondary follicle oocytes from P10 ovaries. The x and y axis of the dot plots are Log_2_CPM from RNA-seq. Fold change > 2, FDR < 0.05. **(B)** Heatmaps showing the DEGs at secondary follicle stage upon NFYA loss, and the NFYA binding, NFYA motif occurrence around the TSS of corresponding genes. Left panel: the downregulated or upregulated genes are separated into two groups based on whether they are affected at primary follicle stage. Right panel: the secondary stage specifically downregulated or upregulated genes are separated into two groups based on whether they have direct NFYA binding. **(C)** Genome browser view of NFYA-targeted infertility related gene examples of NFYA CUT & RUN and RNA-seq of secondary follicles. **(D)** Heatmap showing late-PFA gene (n=834) expression in WT and *Nfya*-cKO oocytes at the indicated stages. **(E)** Heatmaps showing the downregulated late-PFA genes (n=548) at secondary follicle stage upon NFYA loss, and the NFYA binding around the TSS of corresponding genes. The downregulated genes are separated into two groups based on whether they have direct NFYA promoter binding. **(F)** Genome browser view of NFYA targeted late-PFA gene examples of NFYA CUT&RUN and RNA-seq in secondary follicles. **(G)** Density plots showing the average ATAC_FC signals in the promoter regions of secondary follicle-specific DEGs with or without NFYA promoter binding. P values were calculated with two-sided Student’s t-test. **(H)** Genome browser view of examples of the NFYA binding (shaded), ATAC and RNA-seq signals in WT and CKO secondary follicles. **(I)** The percentage of early- and late-PFA genes regulated by NFYA in primary and secondary follicle oocytes.

To determine the role of NFYA in regulating the primary-to-secondary follicle transition, we first focused on the 1,888 early-PFA genes (**Figure 2F**). We found that these genes are activated at primary follicle oocytes and their expression is maintained in secondary follicle oocytes (**Figure S3I**). Of these early-PFA genes, 661 are affected by NFYA deficiency, including 389 affected in the primary follicles and 576 affected in the secondary follicles with 304 commonly affected in both stages (**Figure S3J**). Of the 272 newly affected early-PFA genes, 110 (40.4%) gene promoters are bound by NFYA, including *Pde6d*, *Myh10*, *Rell1*, and *Fam72a* (**Figure S3K**). Notably, PDE6D has been shown to be important for cAMP-PKA and PI3K-AKT signaling, which are critical for PFA ^49,53^, suggesting a critical role of NFYA in regulating PFA. Motif analysis of the 162 genes revealed 82.1% (133/162) contain at least one NFYA binding motif in their promoter regions (**Figure S3L**).

We also identified 834 genes specifically activated during the primary-to-secondary transition by comparing the transcriptomes of primary and secondary follicle oocytes, termed late-PFA genes (**Figure 3D and Table S2**). Of these late-PFA genes, 548 (65.7%) failed to be activated due to NFYA depletion (**Figure 3D**). A global disruption of late-PFA genes expression in *Nfya*-cKO secondary follicle oocytes was further validated by GSEA (**Figure S3M**). Among the 548 affected late-PFA genes, 229 gene promoters are directly bound by NFYA (**Figure 3E**), including *Katnal2* and various histone genes (**Figure 3F**). Motif analysis on the 319 genes revealed 84% (268/319) contain at least one NFYA binding motif in their promoter regions (**Figure S3N**). Collectively, these data suggest a direct role of NFYA in late-PFA regulation.

To explore whether NFYA activates late-PFA genes by promoting chromatin accessibility in secondary follicle oocytes, we performed ATAC-seq in *Nfya*-cKO and WT secondary follicle oocytes (**Figure S3O**). Comparative analysis showed that NFYA deficiency significantly reduced chromatin accessibility at the promoters of NFYA-bound downregulated genes (**Figure S3D, right**). Importantly, promoter accessibility of NFYA-bound secondary follicle oocyte-specific downregulated genes (n=658) also significantly decreased (for example *Hist1h2bk/j*, *Nfe2l1*, *Xbp1*, *Ccnb2*, *Cdc23*, and *Top2a*) (**Figure 3G-H**). Collectively, these data suggest that NFYA is essential for gene activation in secondary follicle oocytes through promoting chromatin accessibility.

In summary, NFYA activates 35.01% of early-PFA genes and 65.71% of late-PFA genes (**Figure 3I**), supporting its role as a critical activator of PFA.

### NFYA loss triggers ferroptosis-like cell death leading to *Nfya*-cKO oocyte degeneration

Next, we asked how NFYA deficiency causes degeneration and elimination of early follicles (**Figure 1K-M**). Previous studies have demonstrated that oocyte elimination can be caused by apoptosis, autophagy, and ferroptosis ^54–56^. Interestingly, GSEA revealed significant enrichment of ferroptosis-related pathways, but not apoptosis-or autophagy-related pathways, in *Nfya*-cKO oocytes compared to WT controls (**Figure 4A and S4A**). Notably, many ferroptosis suppressors, including *Pdia4*, *Hspa5*, *Nfe2l1*, and *G3bp1* ^57–60^, are downregulated in *Nfya*-cKO secondary follicle oocytes and are directly bound by NFYA (**Figure 4B**), suggesting that NFYA protects oocytes from ferroptosis. Consistently, the protein levels of apoptosis-related markers (γH2AX and cleaved caspase3) or autophagy-related marker LC3 are not changed in *Nfya*-cKO oocytes (**Figure S4B-C**). The labile iron ions (Fe^2+^), a key marker of ferroptosis detected by FerroOrange, exhibit a markedly altered cytoplasmic distribution pattern in *Nfya*-cKO secondary follicle oocytes, but not in primary follicle oocytes (**Figure 4C and S4D**). Interestingly, the overall signal intensity of FerroOrange is comparable between *Nfya*-cKO and WT oocytes (**Figure S4E**), diverging from canonical ferroptosis, which typically features increased Fe² levels ^56,61^. Since Fe²-driven lipid peroxidation of cellular membranes is a key step in ferroptosis ^61,62^, we examined lipid peroxidation using the BODIPY 581/591 C11 probe (a lipid peroxidation sensor) and found that *Nfya*-cKO oocytes exhibit significantly increased lipid peroxidation, particularly along the cell membrane (**Figure 4D**). Furthermore, as mitochondrial and ER membranes are highly sensitive to lipid peroxidation ^61,62^, we examined mitochondria and ER membrane structure by electron microscope. The results showed loss of cristae structure and disruption of mitochondrial membranes in *Nfya*-cKO secondary follicle oocytes (**Figure 4E**). We also observed disorganized ER, which stacked together, formed a big circler shape, and contacted with disrupted mitochondria (**Figure 4F-G**), similar to the previously reported ER disorganization induced by strong ER stress ^63,64^. Consistently, NFYA-bound downregulated genes were enriched for ER stress in *Nfya*-cKO oocytes (**Figure S3G**).

**Figure 4.**
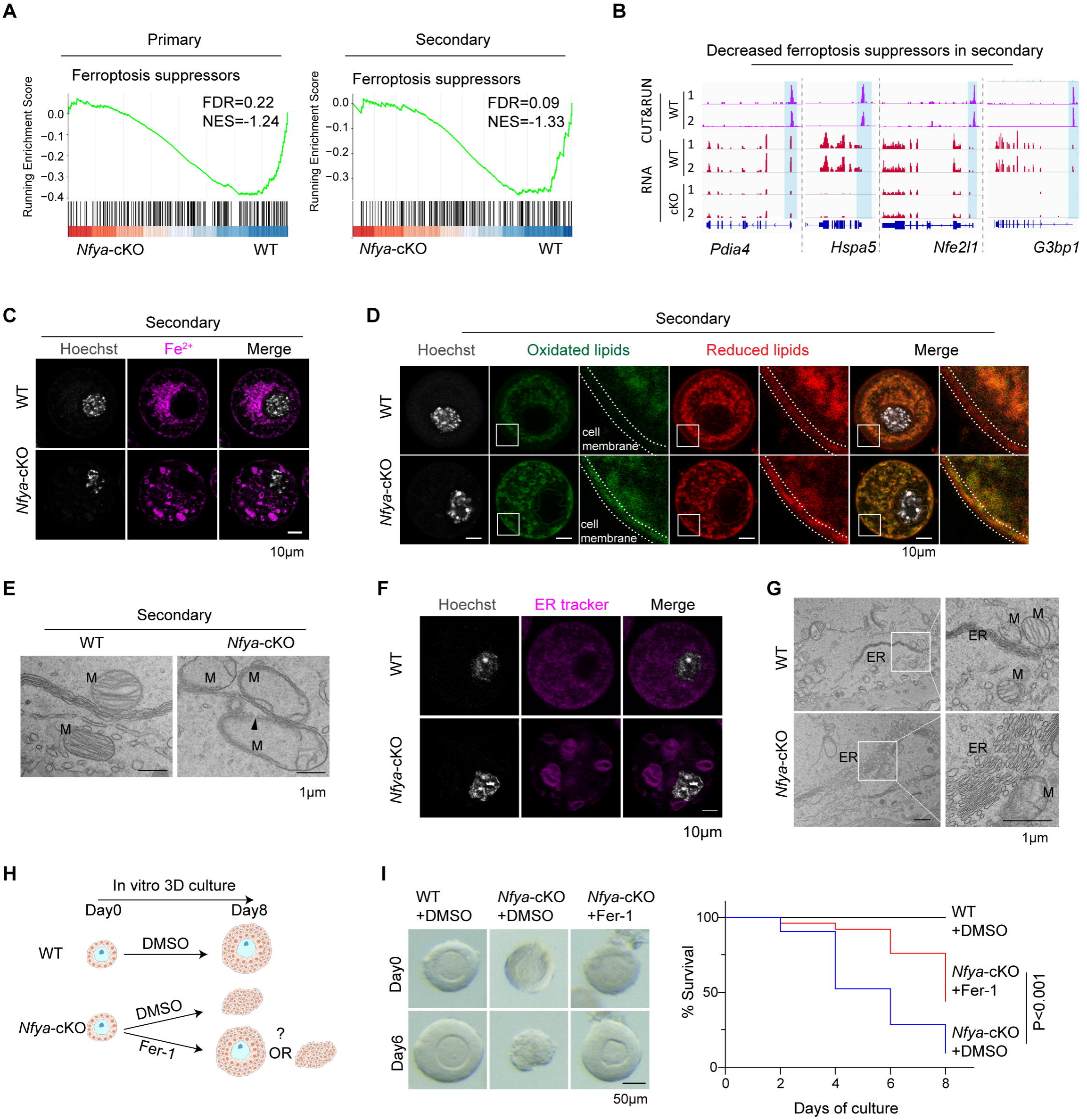
NFYA loss triggers non-canonical ferroptosis in growing oocytes. **(A)** Gene set enrichment analysis (GSEA) revealed that ferroptosis suppressors are significantly decreased in *Nfya*-cKO oocytes. **(B)** Genome browser view of examples of NFYA CUT&RUN and RNA-seq of NFYA-targeted ferroptosis suppressors in secondary follicles. **(C)** Representative images of FerroOrange (an Fe^2+^ indicator) staining of WT and *Nfya*-cKO secondary follicle oocytes reveal a change in staining patterns. Scale bar, 10 μm. **(D)** Representative images of BODIPY 581/591 C11 probe staining of oxidized lipids and reduced lipids in WT and *Nfya*-cKO secondary follicle oocytes. Scale bar, 10 μm. **(E)** Electron Microscopy (EM) images showing the cross-section of mitochondria (M) in WT and *Nfya*-cKO secondary follicle oocytes. The arrowhead indicates damaged mitochondria membrane. Scale bar, 1 μm. **(F)** Representative images of Endoplasmic Reticulum (ER) tracker labeling in WT and *Nfya*-cKO secondary follicle oocytes reveal morphological changes in ER. Scale bar, 10 μm. **(G)** EM images showing the cross-section of ER in WT and *Nfya*-cKO secondary follicle oocytes. Right are the high-magnification EM images of left boxed areas showing the ER-mitochondria contact. Scale bar, 1 μm. **(H)** Schematic diagram of an *in vitro* 3D follicle culture experiment with the isolated WT follicles treated with DMSO and the *Nfya*-cKO follicles treated with DMSO or Fer-1. (**I)** Left panel: representative images of a WT or *Nfya*-cKO follicle at day 0 or day 6 after culturing with DMSO or Fer-1. Scale bar, 50 μm. Right panel: Survival curves of WT and *Nfya*-cKO follicles in culture supplemented with DMSO or Fer-1. P values, log-rank test.

To determine whether ferroptosis plays a causal role in *Nfya*-cKO oocyte degeneration, we cultured follicles in a 3D *in vitro* system for 8 days with or without the ferroptosis antagonist ferrostatin-1 (Fer-1) (**Figure 4H**). We found that the presence of Fer-1 significantly reduced NFYA deficiency-induced oocyte degeneration (**Figure 4I**) supporting a causal role of ferroptosis in *Nfya*-cKO oocyte degeneration. Taken together, these results suggest that NFYA loss triggers ferroptosis-like cell death in oocytes, characterized by decreased expression of ferroptosis suppressors, abnormal cytoplasmic Fe^2+^ distribution, ER disorganization, excessive lipid peroxidation, and rupture of cellular membrane.

### dTAG-based NFYA degradation impairs ZGA

After demonstrating NFYA is a key regulator of PFA, we next asked whether NFYA is also required for ZGA. Although our previous studies have suggested a role in ZGA through siRNA-mediated *Nfya* knockdown in cultured GV oocytes ^24^, the potential for off-target siRNA effects or secondary effects of GV oocyte culture could not be excluded. Because NFYA is required for normal oocyte development, the *Nfya*-cKO model cannot address its role in ZGA; however, the rapid protein degradation dTAG system, recently applied in mouse embryos ^30,31^ makes this possible.

To this end, we designed an N-terminal HA (hemagglutinin)-FKBP (FK506 binding proteins)-NFYA fusion construct (referred to as NFYA^dTAG^) to enable conditional degradation (**Figure S5A**). After confirming the stability of this fusion protein and the efficiency of dTAG13-induced degradation in mouse ES cells (mESCs) (**Figure S5B-C**), we generated the *Nfya*^dTAG^ mouse line using the CRISPR technology ^65^ (**Figure S5D-E**). Notably, these mice are fertile and developmentally normal, with an average little-size of 10 pups from 11 litters obtained from *Nfya*^dTAG/dTAG^ intercrosses (**Figure S5E**), indicating that the dTAG knock-in does not impair NFYA function under physiological conditions. We then confirmed that dTAG13 treatment can efficiently deplete NFYA from cultured embryos at late 2-cell (L2C) stage (**Figure 5A**). Collectively, these results demonstrate the successful generation of a NFYA^dTAG^ mouse model.

**Figure 5.**
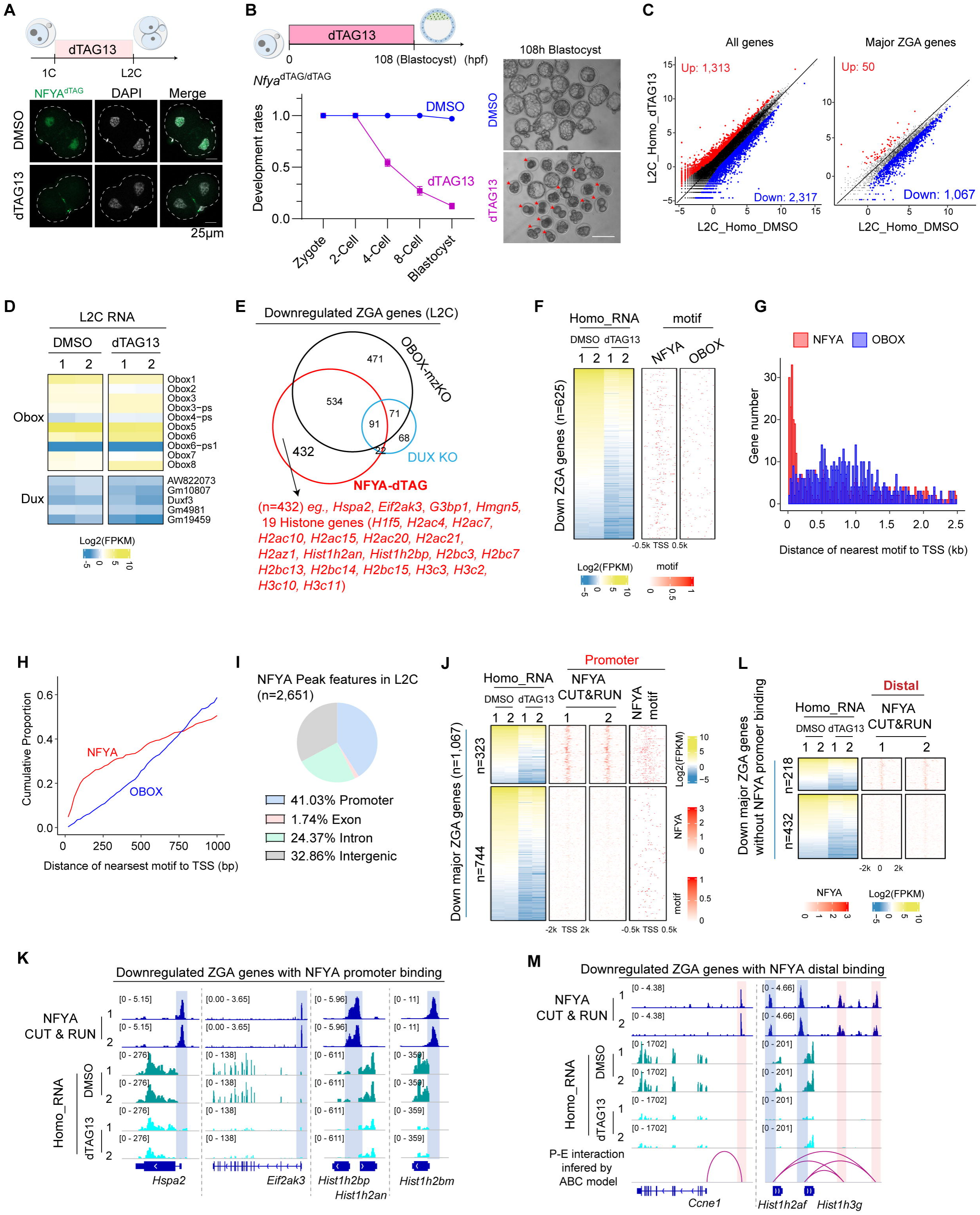
dTAG13-mediated NFYA depletion causes ZGA defects. **(A)** Immunostaining confirming dTAG13-induced NFYA degradation during ZGA. Anti-HA antibody was used for endogenous NFYA^dTAG^ fusion protein detection. Scale bar, 25 μm. **(B)** Left upper panel: schematic diagram of the experimental design. Left lower panel: The *Nfya*^dTAG/dTAG^ (Homo) embryo development rate upon DMSO or dTAG13 treatment. The data are presented as mean ± SEM. Right panel: representative images of blastocyst stage embryos after NFYA degradation (DMSO, n = 83; dTAG13, n = 90; n represents the total number of embryos from three independent experiments). Scale bar, 100 μm. Red arrows indicate representative arrested embryos in dTAG13 group. **(C)** Scatter plots comparing the gene expression profiles of Homo late-2cell embryos treated with dTAG13 or DMSO. The x and y axis of the dot plots are Log_2_CPM from RNA-seq. Fold change > 2, FDR < 0.05. A total of 2,773 major ZGA genes are analyzed (Table S2). **(D)** Heatmaps showing the *Obox* and *Dux* family gene expression in late 2-cell embryos treated with dTAG13 or DMSO. **(E)** Venn diagram showing the overlapped downregulated ZGA genes in L2C embryos among *Obox* mzKO ^39^, *Dux* knockout ^75^, and NFYA depletion (NFYA-dTAG). **(F)** Heatmaps showing the commonly downregulated ZGA genes (n=625) in OBOX-mzKO and NFYA-dTAG, and the motif occurrence of NFYA and OBOX around the TSS of the corresponding genes. **(G)** Bar chart showing the downregulated gene numbers based on the distance from their TSS to nearest NFYA or OBOX motif. **(H)** Cumulative distributions of the distance from commonly downregulated gene TSS to the nearest NFYA or OBOX motif. **(I)** The genomic distribution of NFYA binding peaks generated by NFYA CUT&RUN using late 2-cell embryos. **(J)** Heatmaps showing the downregulated major ZGA genes (n=1,067) at late 2-cell embryos upon NFYA loss, and the NFYA binding, NFYA motif occurrence around the TSS of corresponding genes. The downregulated major ZGA genes are separated into two groups based on whether they have direct NFYA promoter binding. **(K)** Genome browser view examples of NFYA CUT&RUN and RNA-seq in NFYA promoter targeted ZGA genes at late-2cell stage. **(L)** Heatmaps showing the downregulated major ZGA genes (without NFYA promoter binding) with predicted P-E interactions (n=650) at late 2-cell stage upon NFYA loss, and the NFYA binding around the center of enhancers. The downregulated major ZGA genes are separated into two groups based on whether their putative enhancers have direct NFYA binding. **(M)** Genome browser view examples of NFYA CUT&RUN and RNA-seq in NFYA distal targeted ZGA genes at late-2cell stage.

To evaluate the role of NFYA in ZGA, we treated *Nfya*^dTAG/dTAG^ (Homo) embryos with dTAG13 and found that NFYA-depletion caused half of the embryos to arrest at 2-3 cell stage and ∼90% to failed to reach blastocyst stage (**Figure 5B and S5F**), suggesting a critical role of NFYA in ZGA and preimplantation development. To directly address the role of NFYA in ZGA, we collected dTAG13-treated Homo late 2-cell (L2C) embryos and performed RNA-seq (**Figure S5G**), which revealed 2,317 downregulated and 1,313 upregulated genes relative to that of the control (**Figure 5C, left and Table S4**). Among the downregulated genes, 1,067 are major ZGA genes (**Figure 5C, right and Table S4**), indicating that NFYA activates these ZGA genes. NFYA degradation did not affect the expression of OBOX or DUX gene families (**Figure 5D**), nor *Nr5a2* or *Klf17* (**Figure S5H**), suggesting that NFYA regulates ZGA independently of these known ZGA regulators. Integrative analysis of DEGs showed that 40% of NFYA-regulated ZGA genes (432 out of 1,067) do not overlap with those of OBOX-or DUX-regulated ZGA genes, including *Hspa2*, *Eif2ak3*, *G3bp1*, and various histone genes (**Figure 5E**). Intriguingly, although we found that 625 major ZGA genes were regulated by both NFYA and OBOX (**Figure 5E-F**), motif analysis revealed distinct binding patterns with NFYA motifs mainly enriched near their core promoter regions (TSS ± 500bp), while OBOX motifs are dispersed throughout the gene body (**Figure 5G-H**), suggesting that NFYA most likely regulates these genes through promoter binding and OBOX may through enhancer binding. Collectively, these data demonstrate that NFYA is critical for ZGA and preimplantation development.

### NFYA activates ZGA genes by promoter and distal binding

To determine whether NFYA directly binds to and regulates major ZGA genes, we performed low-input CUT&RUN for NFYA in L2C embryos. For quality control purpose, we confirmed the reproducibility and compared our low-input L2C dataset with that of bulk mESC dataset and found most of the major peaks are similar (**Figure S6A-C**). The most enriched motif in L2C peaks is the classical NFY binding sequence CCAAT (**Figure S6D**). More than half of the NFYA binding peaks are located in distal regions (putative enhancers) (**Figure 5I**), which is different from the binding patterns in oocytes (**Figure S2E and S3B**) or mESCs (**Figure S6E**) where it mostly binds to promoters. To verify that the distinct NFYA-binding patterns observed in L2C embryo are genuine, we performed NFYA CUT&RUN in NFYA-depleted L2C embryos and found that both promoter and distal NFYA-binding peaks disappeared in NFYA-depleted L2C embryos (**Figure S6F**), suggesting that NFYA indeed exhibits preferential binding to distal regions in L2C. To investigate whether NFYA-bound distal regions are enriched for transposable elements, we performed analysis which revealed that NFYA-bound sites are preferentially associated with LTR, SINEs, and snRNA repetitive elements (**Figure S6G**). Further analysis showed that NFYA binding is significantly more enriched on downregulated ZGA genes compared to other downregulated genes (**Figure S6H**), suggesting that NFYA preferentially binds and regulates ZGA genes. Integrative analyses of CUT&RUN and RNA-seq data revealed that many downregulated major ZGA genes (323 out of 1,067) have promoter NFYA binding, including *Hspa2*, *Eif2ak3*, *G3bp1*, *Hmgn5*, and various histone genes (**Figure 5J-K and S6I**). Further motif analysis of the 744 down-regulated genes without CUT&RUN signal revealed 72% (538/744) contain at least one NFYA motif in their promoter regions (**Figure S6J**). Collectively, these data suggest a direct role of NFYA-promoter binding in ZGA regulation.

To further characterize NFYA distal binding, we utilized the activity-by-contact (ABC) model ^66^ to predict potential NFYA-mediated promoter-enhancer (P-E) interactions and found that 650 out of 744 downregulated ZGA genes are correlated with predictable enhancer activity (**Figure S6K**). Further analysis indicated that many NFYA-bound enhancers are linked to downregulated major ZGA genes (218 out of 650), including *Ccne1*, *Hist1h2af*, *Sael*, and *Polr3g* (**Figure 5L-M and S6L**). Collectively, our integrated analysis supports that NFYA directly regulates ZGA genes by both promoter and distal binding.

To explore whether NFYA regulates ZGA genes by promoting chromatin accessibility in L2C, we performed ATAC-seq in L2C embryos with or without dTAG13 treatment (**Figure S6M**). Comparative analysis showed that NFYA depletion reduced chromatin accessibility at the promoters of downregulated ZGA genes (**Figure S6N**), but not the overall NFYA distal binding (**Figure S6O**), suggesting that NFYA contributes to gene activation during ZGA by promoting chromatin accessibility.

### NFYA regulates PFA and ZGA by multifaceted chromatin binding

Having demonstrated that NFYA serves as a common regulator for both PFA and ZGA through its promoter and distal chromatin binding, we next sought to examine the conserved and distinct roles of NFYA in these two major genome transcriptional activation events. By comparing NFYA binding profiles in oocytes and L2C embryos, we found that 60.9% oocyte-specific NFYA binding is located at promoter regions, whereas 85.5% L2C-specific NFYA binding is located at distal regions (**Figure 6A**), underscoring the distinct preference of NFYA-chromatin interactions during PFA and ZGA. GO analysis revealed that oocyte-specific NFYA bound genes were associated with “response to hormone stimulus” and “meiotic cell cycle”, whereas L2C-specific NFYA bound genes were associated with “stem cell maintenance” and “RNA splicing” (**Figure S7A**), suggesting distinct functions of stage-specific NFYA-chromatin interactions. Integrating CUT&RUN and ATAC-seq revealed that the L2C-specific NFYA binding sites are largely inaccessible in oocytes (**Figure S7B**), suggesting that stage-specific chromatin accessibility may contribute to the differential NFYA binding landscape. Additionally, motif enrichment analysis of the NFYA oocyte-and L2C-specific binding regions revealed that, in addition to the enrichment of NFYA motif, TFs such as SP1, KLF15, and CTCF are enriched at oocyte-specific NFYA-binding regions, whereas NR5A2, RARG, and OTX2 are enriched at L2C-specific regions (**Figure 6B**), suggesting that these stage-specific factors may help shape the multifaceted NFYA chromatin binding profiles during PFA and ZGA.

**Figure 6.**
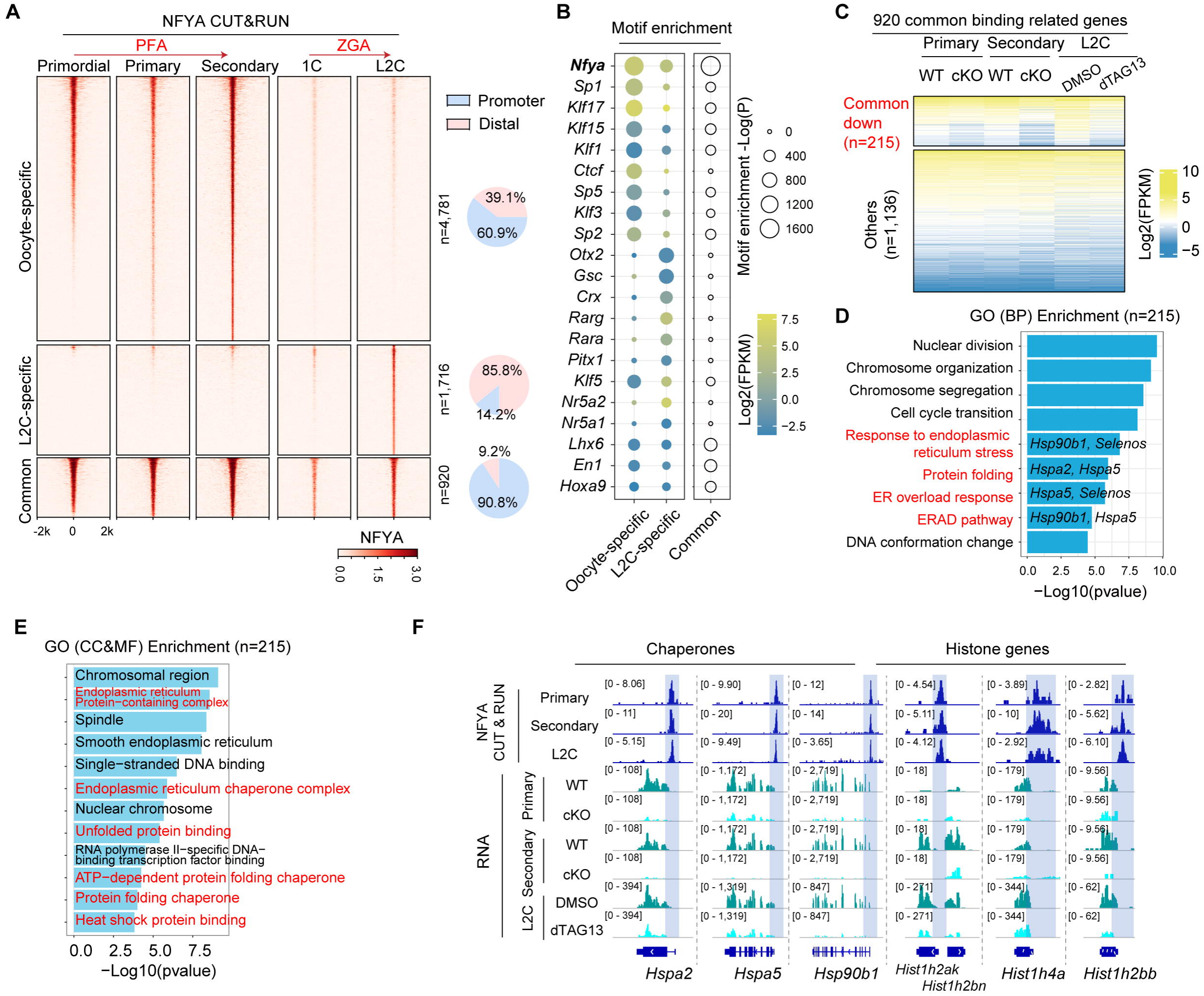
NFYA regulates PFA and ZGA by multifaceted chromatin binding. **(A)** Heatmaps showing the dynamics of NFYA genomic binding profile during PFA (primordial follicle oocytes, primary follicle oocytes, and secondary follicle oocytes) and ZGA (zygotes and late 2-cell embryo) at promoter and distal regions. Pie chart showing the percentage of promoter and distal occupancy. **(B)** Enrichment and expression of top TF motifs at the CUT&RUN peak groups (oocyte-specific, L2C-specific, and common). **(C)** Heatmaps showing the gene expression of the 920 common binding associated genes (n=1,351) in primary follicle oocytes, secondary follicle oocytes, and L2C embryo upon NFYA loss. These genes are separated into two groups: commonly downregulated genes in both oocytes and L2C (n=215) and other genes (n=1,136). **(D)** Representative GO (BP) terms enriched in the commonly downregulated genes (n=215). The example genes of the highlighted term are shown. **(E)** Representative GO (CC and MF) terms enriched in the commonly downregulated genes (n=215). **(F)** Genome browser view of NFYA CUT&RUN and RNA-seq of example NFYA target genes common in oocytes and L2C.

### Pre-occupied NFYA regulates chaperones and histone genes common in both PFA and ZGA

In addition to the stage-specific binding, we also identified 920 common NFYA binding regions, predominantly (90.8%) located at promoter regions (**Figure 6A**). Intriguingly, the 920 common bound regions not only showed stronger NFYA binding signals and motif enrichment (**Figure 6A-B**), but also were already pre-occupied in both the primordial follicle stage and the zygotic stage before PFA or ZGA begins, whereas most of the oocyte-or L2C-specific NFYA bound regions were gradually established during transcriptional activation (**Figure 6A and S7C**). Consistent with the pre-occupancy, NFYA immunostaining also showed nuclear enrichment at primordial follicle stage and zygotic stage (**Figure 1F**). Consequently, these 920 common NFYA bound regions were accessible before PFA or ZGA begins (**Figure S7B**), and NFYA-depletion reduced their accessibility in both oocytes and L2C (**Figure S7D**), suggesting that NFYA pre-occupancy is important for the chromatin opening at those regions. Intriguingly, we found that chromatin accessibility was reduced more at the common bound regions compared with the oocyte-or L2C-specific bound regions (**Figure S7E**), suggesting that NFYA is more important for maintaining chromatin accessibility at the commonly bound regions than at stage-specific bound regions. In sum, these findings suggest that NFYA plays a more important role in the common and pre-occupied regions.

Integrating CUT&RUN and RNA-seq, we identified 1,351 genes associated with the 920 common pre-bound regions. Among them, 215 are commonly downregulated in both PFA and ZGA upon NFYA-depletion (**Figure 6C**). GO analysis revealed that they are associated with Biological Processes, such as nuclear organization, ER response, and protein folding (**Figure 6D**); Cellular Components, such as chromosomal region and endoplasmic reticulum; and Molecular Functions, such as ER chaperone complex and heat shock protein binding (**Figure 6E**), including chaperones and histone genes (**Figure 6F**). Collectively, these results suggest that NFYA regulates both PFA and ZGA through a combination of divergent and conserved chromatin binding. Additionally, pre-occupied NFYA regulates chaperones and histone genes common in both PFA and ZGA.

### Inhibition of chaperones impairs both PFA and ZGA

Histone genes contribute to genome activation by affecting nuclear organization ^67^, but the role of chaperones in PFA or ZGA remains unclear. To investigate the function of NFYA-regulated chaperones in PFA and ZGA, we used the chaperone inhibitors Ganetespib (1 nM∼150 nM for cells) ^68,69^ and VER155008 (1 μM∼50 μM for cells) ^70^ that respectively inhibit HSP90 or HSPA2/5. Titration in mouse embryos (2 nM, 5 nM, 10 nM, and 25 nM Ganetespib; 1 μM, 5 μM, 10 μM, and 25 μM VER155008) (**Figure S7F**) showed that 10 nM Ganetespib or 10 μM VER155008 caused 2-cell embryo arrest, while 5 nM Ganetespib and 5 μM VER155008 had only a minor effect on blastocysts development. Since NFYA-depletion downregulated both HSPA2/5 and HSP90, and to avoid potential off-target effects caused by high concentration inhibitor treatment, we performed the treatment by combining 5 nM Ganetespib and 5 μM VER155008 (referred to as G&V hereafter). Only treatment before ZGA (G&V 1C-B), but not after ZGA (G&V 4C-B), arrest most embryos at the 2-4 cell stage (**Figure 7A**), suggesting a specific role of chaperones in ZGA. To directly address the role of chaperones in ZGA, we collected G&V-treated L2C embryos and performed RNA-seq (**Figure S7G**), which revealed 705 downregulated and 718 upregulated genes compared to that of the control (**Figure 7B, left and Table S5**). Among the downregulated genes, 181 are major ZGA genes (**Figure 7B, right and Table S5**), indicating that chaperones have a role in activating these ZGA genes. Moreover, half of the downregulated ZGA genes were overlapped with NFYA-activated ZGA genes, including cell fate regulator: *Tead4* and *Fgf4*; and RNA processing factors: *Ddx49*, *Rc3h2*, and *Mov10l1* (**Figure 7C),** suggesting that chaperones contribute to ZGA. Collectively, these data suggest that chaperones may play an important role in ZGA and preimplantation development.

**Figure 7.**
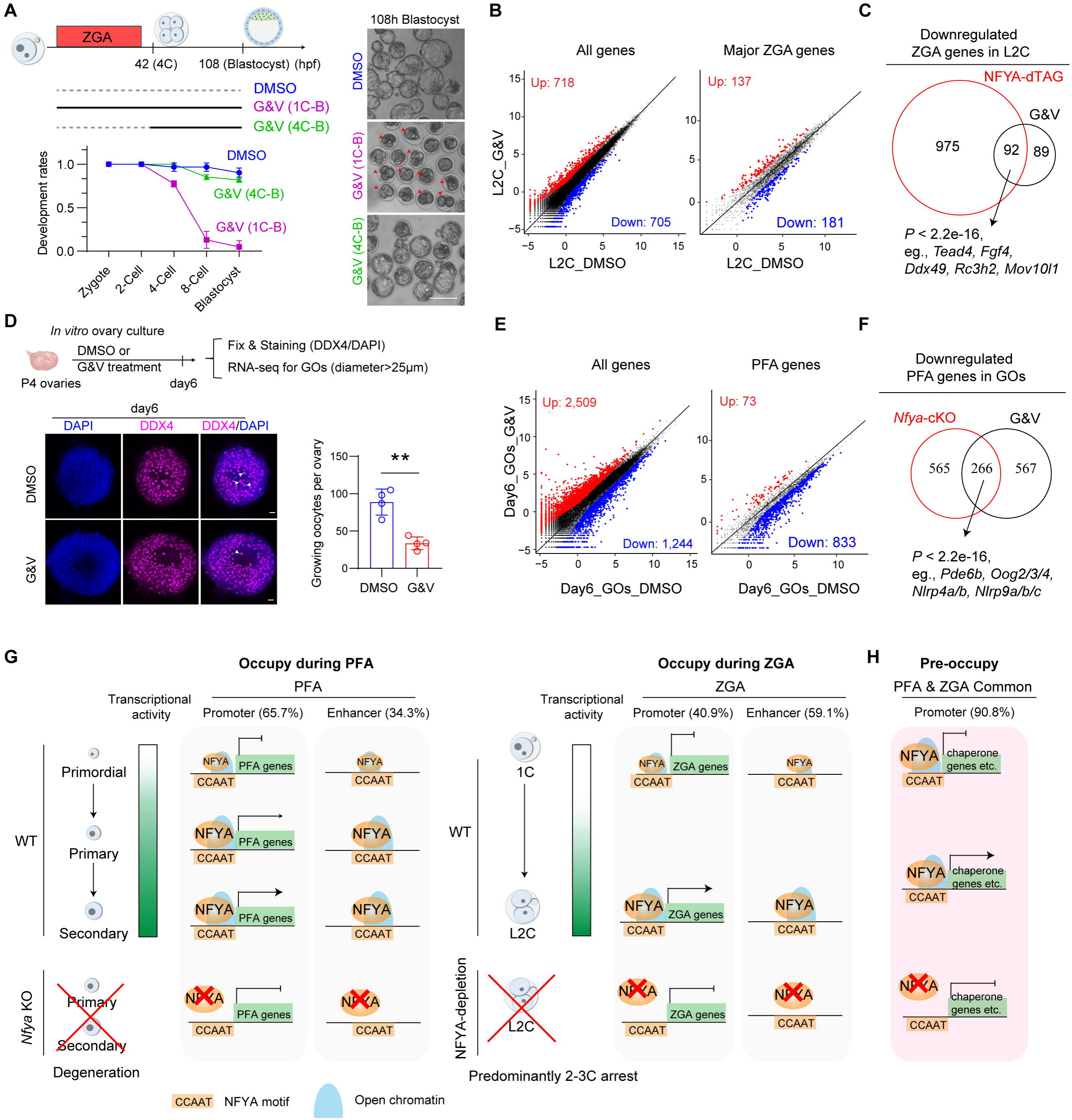
Inhibition of chaperones impairs both PFA and ZGA. **(A)** Left upper panel: schematic diagram of the experimental design. Left lower panel: the embryo development rate upon DMSO or G&V treatment (Ganetespib 5nM and VER155008 5μM). The data are presented as mean ± SEM. [DMSO, n = 67; G&V (1C-B), n = 64; G&V (4C-B), n = 47. n represents the total number of embryos from two independent experiments]. Right panel: representative images of blastocyst stage embryos after chaperones inhibition (G&V). Scale bar, 100 μm. Red arrows indicate arrested embryos in G&V (1C-B) group. **(B)** Scatter plots comparing gene expression profiles of late-2cell embryos treated with G&V or DMSO. The x and y axis of the dot plots are Log_2_CPM from RNA-seq. Fold change > 2, FDR < 0.05. ZGA genes, n=2,773. **(C)** Venn diagram showing the overlapped downregulated ZGA genes between NFYA depleted (NFYA-dTAG) and G&V treated L2C embryos. P values (Fisher’s exact test, two-sided) for overlapped genes are also shown. **(D)** Upper panel: schematic diagram of the experimental design. Lower left panel: ovarian morphology of 6 days *in vitro* cultured P4 ovaries with DMSO and G&V treatment. DDX4 stained oocytes. The white arrowhead indicates representative growing oocytes. Scale bar, 50 μm. Lower right panel: numbers of growing oocytes (DDX4 positive oocytes bigger than 25 μm in diameter that were surrounded by cuboidal granulosa cells) in each ovary after 6 days culture with DMSO and G&V treatment. Quantitative data are shown as mean ± SEM; ∗∗p < 0.01, Student’s t test. **(E)** Scatter plots comparing the gene expression profiles of growing oocytes (GOs) from 6 days *in vitro* cultured P4 ovaries with G&V or DMSO. The x and y axis of the dot plots are Log_2_CPM from RNA-seq. Fold change > 2, FDR < 0.05. PFA genes, n=2,463. **(F)** Venn diagram showing the overlapped downregulated PFA genes between *Nfy*a-cKO and G&V treated GOs. P values (Fisher’s exact test, two-sided) for overlapped genes are also shown. **(G)** Model illustrating the role of NFYA in PFA and ZGA. Before PFA (primordial), the PFA genes are silenced. Subsequently (primary and secondary), NFYA binding promotes open chromatin and activates PFA genes through promoter and distal binding. Loss of NFYA leads to reduced chromatin accessibility, defective PFA, and early follicular degeneration. Before ZGA (1C), the ZGA genes are silenced. Subsequently (L2C), NFYA activates ZGA genes through promoter and enhancer binding. Loss of NFYA leads to defective ZGA and predominantly embryo arrest at 2-3 cell stage. **(H)** NFYA pre-occupies and regulates a set of genes, including chaperones and histone genes, common in both PFA and ZGA through conserved promoter binding.

Similarly, to test chaperones’ role in PFA, we treated P4 neonatal ovaries, when PFA begins in vivo, with G&V and found a significant reduction by day 6 in growing oocytes (GOs), defined as DDX4-positive oocytes larger than 25 μm surrounded by cuboidal granulosa cells (**Figure 7D**), suggesting an important role of chaperones in PFA. Notably, defects were more severe at day10 (**Figure S7H**), with significant loss of central growing follicles, suggesting defective PFA and oocyte arrest. RNA-seq of growing oocytes from G&V-treated (6 day) ovaries (**Figure S7I**) revealed 1,244 downregulated genes relative to that of the control (**Figure 7E, left and Table S6**). Among the downregulated genes, 833 are PFA genes (**Figure 7E, right and Table S6**), indicating that chaperones have an important role in activating these PFA genes. Moreover, 266 of the 833 downregulated PFA genes overlapped with NFYA-activated PFA genes, including Oog2/3/4 and NLRP family (**Figure 7F),** suggesting that chaperones may contribute to the regulation of these PFA genes. Taken together, these data suggest that chaperones may be critical for both PFA and ZGA.

Collectively, our data support a common model for NFYA’s function in transcriptional activation during PFA and ZGA: 1) NFYA pre-occupies and regulates a group of common genes including those encoding chaperones and histones before genome activation; and 2) NFYA gradually establishes binding at oocyte-or L2C-specific genes to activate them during the two genome activation events.

## Discussion

Although PFA and ZGA share many molecular hallmarks ^8,71^, a common regulator has not been identified. Here, we demonstrate that NFYA is the long sought-after TF involved in both events (**Figure 7G-H**). NFYA exhibits distinct chromatin-binding preferences with predominant promoter targeting during PFA and enhancers targeting during ZGA, thereby regulating transcriptional programs in a cell type-and developmental stage-specific manner. On the other hand, conserved NFYA-chromatin binding pre-occupies and regulates a set of common genes, including chaperones and histone genes, in both PFA and ZGA. Together, NFYA is a common and multifaced regulator of PFA and ZGA.

### Role of NFYA in quiescent primordial follicle activation

A major function of ovary is to balance the quiescent and activated primordial follicle pool. The of primordial follicles quiescence is maintenaned by a regulatory network, including FOXO3A ^10^, LHX8 ^12,14^, and PTEN-PI3K signaling pathway ^9^, whose knockout causes excessive activation of PFA genes and subsequent depletion of primordial follicles. However, the pioneer factors that directly activate PFA *in vivo* remain largely unknown. Although NOBOX has been reported to be critical for PPT ^72^, extensive transcriptional defects and dysregulation of early germ cell genes have been observed in oocytes from newborn ovaries before primordial follicle formation ^73^, suggesting that the defective PPT might be a secondary effect of earlier germ cell abnormalities. Here, we show that NFYA directly activates a subset of PFA genes through direct promoter binding (**Figure 2-3**) and that its oocyte-specific loss depletes primary and secondary follicles (**Figure 1**), supporting a critical role of NFYA in PFA. This raises several mechanistic questions, including how NFYA deals with the known regulators to drive primordial follicles out quiescence. Intriguingly, an interaction between NFYA and FOXO3A has been reported in other cell types ^74^, suggesting that they may coordinately regulate the balance between quiescence and activation of primordial follicles. Characterizing their common targets during this transition may clarify how this balance is controlled at the molecularly level.

### Role of NFYA in ZGA

Several TFs important for mammalian ZGA have been identified, including OBOX ^39^, DUX ^75^, NR5A2 ^41,43^, GABPA^31^, and KLF17 ^76^, although none of them has a demonstrated function in PFA. Although our previous studies implicated NFYA in ZGA ^24^, definitive evidence was lacking. Here we provide three lines of evidence for its role in ZGA. First, dTAG-mediated NFYA degradation downregulated 1,067 ZGA genes (**Figure 5C**), a number similar to that affected by OBOX ^39^. Second, many of the down-regulated genes are direct targets with NFYA bound to their promoters or distal regions (**Figure 5J-M and S6I-L**). Third, acute NFYA depletion before ZGA arrests half of the embryos at 2-3 cell stage, consistent with dysregulation of ZGA.

Cell type-or stage-specific TFs have advanced our understanding of the intricate gene regulatory networks, but conserved mechanisms underlying similar transcriptional programs across different contexts are often overlooked. The conserved NFYA-chromatin interactions for activating chaperone and nucleosome organization related genes that we uncovered underscore the importance of these processes in both PFA and ZGA (**Figure 6-7**). Intriguingly, it was recently demonstrated that successful ZGA is closely associated with proper nucleosome organization ^67^. Although chaperones are reported as the first major products of mouse ZGA ^77^, their roles in oocytes and early preimplantation development remain unclear. Our inhibition experiments suggest that chaperones may be critical for both PFA and ZGA, and the 2-4 cell stage arrest caused by inhibitor treatment before but not after ZGA supports a ZGA-specific requirement rather than a general toxicity. However, how the chaperones affect ZGA remains to be determined. In an indirect model, chaperones maintain the stability and activity of the proteins that execute ZGA (such as transcription factors, chromatin regulators and epigenetic factors) and buffer the folding load created by maternal protein turnover and the surge in new translation, with reduced ZGA gene expression as a downstream consequence. A more direct model would involve the chaperones acting on nuclear factors that regulate transcription, for example by promoting the engagement of paused RNA polymerase II, although these mechanisms are speculative. Given that comprehensive epigenetic reprogramming accompanies both PFA and ZGA ^8,78^, the interplay between NFYA-chromatin binding and epigenetic reprogramming warrants investigation. Since NFYA is evolutionarily conserved in all eukaryotes ^79^, whether it has evolutionarily conserved roles in ZGA or PFA in other species remains to be shown.

### NFYA activates transcription programs across diverse biological contexts

Beyond PFA and ZGA, transcription reactivates after mitosis (post-mitotic reactivation), restoring gene expression in the highly compacted and transcriptionally quiescent metaphase genome ^80^. NFYA has been implicated in post-mitotic reactivation ^81^; together with its role in PFA and ZGA, NFYA appears to be a general regulator of transcriptional reactivation across diverse cellular contexts. Activation of dormant embryos also involves genome-wide transcriptional activation ^82,83^, but a role for NFYA in this process is unknown. Activation of quiescent hematopoietic stem cells (HSCs) and tissue regeneration involve less genome-wide transcriptional activation than PFA and ZGA, but also involve transcriptional reprogramming, cell cycle transitions, and rapid protein synthesis and folding ^84,85^. Notably, NFYA also regulates these regenerative processes ^86,87^, suggesting a broader role in reactivating transcription programs. While loss of NFYA in various biological processes has been associated with the induction of apoptosis ^64,86,87^, our results showed that NFYA deficiency in oocytes triggers ferroptosis-like death instead of apoptosis. This divergence underscores the context-dependent roles of NFYA in regulating cell death decisions.

### Limitations of the study

Using NFYA-dTAG system, we found that NFYA-depletion caused ∼50% embryo arrest at 2-3 cell stage, while ∼40% embryo arrest between 4C to morula. Although the early arrest phenotype is within a similar range to those previously established ZGA regulators: OBOX mzKO at hcG-100h showed around 22.2% 4C, 16.7% 3C, 16.7% 2C, and 22.2% 1C embryo arrest, with an additional 22.2% degenerated embryos, while KLF17-mKO showed around 20%∼40% 2C embryo arrest at 96hpi), the later embryonic arrest observed following NFYA depletion could also reflect an essential requirement for NFYA at or after the 4-cell stage. Stage-specific NFYA degradation will therefore be necessary to distinguish its stage-specific requirements and functions of NFYA during early embryonic development.

Although more than half of NFYA-binding peaks in L2C embryos are located in distal regions (**Figures 5I and S6F**), NFYA depletion did not significantly change overall distal chromatin accessibility, in contrast to its effect at promoters. This raises several possibilities: 1) NFYA may promote chromatin accessibility only at a subset of distal regions; 2) NFYA may regulate distal regions through histone modifications rather than directly altering chromatin accessibility; 3) NFYA may be recruited to distal regions by other pioneer factors rather than functioning as a pioneer factor itself.

NFYA loss triggers ferroptosis-like cell death in oocytes, which is characterized by decreased expression of ferroptosis suppressors, redistribution of cytoplasmic Fe2+, excessive lipid peroxidation, ER disorganization, and mitochondria damage. NFYA deficient oocytes could serve as a good model of non-canonical ferroptosis, featuring redistribution rather than elevation of cytoplasmic Fe2+ (**Figure 4C and S4D-E**). Although many ferroptosis suppressors are downregulated by NFYA loss, the mechanisms by which NFYA loss leads to redistribution of cytoplasmic Fe2+ and ER disorganization, as well as how altered Fe2+ distribution initiates ferroptosis, require further investigation.

While our analysis has been mainly focused on the down-regulated NFYA direct targets, we also observed NFYA binding on many upregulated genes in *Nfya*-cKO oocytes (**Figure 2C and 3B**), indicating that NFYA could also have transcriptional repressor function. Further investigations are needed to understand how NFYA functions as a transcriptional repressor in oocytes.

Although we used milder chaperone inhibitor concentrations (Figure 7A-F), potential off-target effects cannot be ruled out, and genetic manipulation studies are needed to understand how chaperones regulate both PFA and ZGA.

## Resource availability

### Lead contact

Further information and requests for resources and reagents should be directed to and will be fulfilled by the lead contact, Yi Zhang

### Materials availability

All unique/stable reagents generated in this study are available from the lead contact with a completed materials transfer agreement.

## Data and code availability

Sequencing data that support the findings of this study have been deposited in the Gene Expression Omnibus (GEO) under accession code GSE295086.

This paper does not report original code.

All other data supporting the findings of this study are available from the corresponding author on reasonable request.

## Acknowledgements

We thank members of the Zhang lab and Qingji Lyu for discussion during the study; Chengjie Zhou and Yota Hagihara for commenting on the manuscript. We thank Nobuyuki Nukina for providing the *Nfya*-flox mouse line; Rodent Histopathology Core for help with HE staining; HMS Electron Microscopy Core for help with electron microscopic imaging; Meng Wang for initial bioinformatic analysis and helpful comments. This project was supported by NIH (R01HD116750) and the HHMI. YZ is an investigator of the Howard Hughes Medical Institute.

## Author contributions

Y.Z. conceived and supervised the project. Q-Y.Y. established the NFYA-dTAG mice and mESCs, performed most experiments including RNA-seq, CUT&RUN, ATAC-seq. B-Y.W. performed the ferroptosis-related analyses and helped in oocytes collection. S.J. performed all the bioinformatic analysis. Q-Y.Y., S.J., B-Y.W., and Y.Z. wrote the manuscript. All authors interpreted the data.

## STAR Methods

### Experimental Model and Study Participant Details

#### Animals

All animal experiments were performed in accordance with the protocols of the Institutional Animal Care and Use Committee at Harvard Medical School. All mice were kept under specific pathogen-free conditions within an environment controlled for temperature (20-22°C) and humidity (40-70%), and were subjected to a 12 h light/dark cycle. *Nfya*^flox/flox^ mice were obtained from Nobuyuki Nukina lab ^64^. *Gdf9-Cre* [Tg(Gdf9-icre)5092Coo/J] mouse lines were purchased from the Jackson Laboratory (011062). The *Gdf9-Cre* mice were crossed with *Nfya*^flox/flox^ mice to generate *Gdf9-Cre Nfya*^flox/flox^ (*Nfya*-cKO) mice. All mice had a C57BL/6J genetic background. Primers used for genotyping are listed in **Table S7**.

*Nfya*-dTAG knock-in mice were generated following a previous protocol with modifications ^31^. Briefly, 2-cell embryo (20hpf) were injected with *Nfya* donor DNA (30 ng/μl), Cas9 protein (25 ng/μl) and sgRNA (50 ng/μl each) using a Piezo impact-driven micromanipulator (Primer Tech, Ibaraki, Japan). After 2 h KSOM incubation, 2-cell embryos were transferred into oviducts of pseudo-pregnant ICR strain mothers (Charles River). F0 chimera mice was backcross with wild-type CD1 mice for at least two generation. Genotyping was performed with mouse tail lysed in lysis buffer (50 mM Tirs-HCl, 0.5% Triton, 400 μg/ml Proteinase K) at 55°C overnight. For F0 and F1 mice genotyping, the primers outside the homology arm are used. For F2 and beyond mice genotyping, the inner primers are used. The primers are included in **Table S7**.

Droplet Digital PCR (ddPCR) was used for detecting the copy number of *Nfya*-dTAG knock-in allele in F1 mice. Briefly, 250 ng purified DNA template was digested by incubation with Haelll Enzyme (NEB) at 37°C for 1h, and then inactivated at 80°C for 5 mins. A final 30 ng DNA was used as template for PCR. *Fkbp* was used for knock-in detecting, *mRPP30* was used as internal control of two copy genome. The PCR primers are listed in Table S7. The mice with single *Fkbp* copy were used for subsequent mating (**Table S8**).

### Cell line, primary culture and establishment of *Nfya*^dTAG^ mESC cell line

Undifferentiated E14 male mouse embryonic stem cells (mESCs) were cultured on 0.1% gelatin coated plates with 2i and LIF condition. E14 mESCs were grown in DMEM (Gibco, 11960069), supplemented with 15% fetal bovine serum (Sigma-Aldrich, F6178), 2 mM GlutaMAX (Gibco, 35050061), 1 mM sodium pyruvate (Gibco, 11360), 1x MEM NEAA (Gibco, 11140050), 0.084 mM 2-mercaptoethanol (Gibco, 21985023), 1 mM sodium pyruvate (Gibco, 11360070), 100 U/ml penicillin-streptomycin (Gibco, 15140122), 1000IU/ml LIF (Millipore, ESG1107), 0.5 μM PD0325901 (Tocris, 4192), and 3 μM CHIR99021 (Tocris, 4423). To establish *Nfya*^dTAG^ mESC cell line, *Nfya* donor DNA and *Nfya*-sgRNA inserted pX330.puro (addgene, #110403) plasmids were co-transfected with Lipofectamine™ 2000 (Invitrogen, 11668030). Cells were then selected with puromycin (Gibco, A1113803), and the single cell-derived colonies were then picked for genotyping and further analysis. Genotyping primers are listed in **Table S7**. Cells were cultured in an incubator containing 5% CO_2_ at 37 °C.

## Method Details

### Preparation and collection of mouse oocytes and embryos

All experiments involved in oocytes and embryo preparation were performed as previously described ^31,35^, with minor modifications. Briefly, primary and secondary follicle oocytes were isolated from mice at postnatal day 7 or day 10. To capture the earlier molecular changes that lead to growing oocytes defects, we collected primordial and primary follicle oocytes at postnatal day 7, when the primary follicle oocytes exhibit no morphological difference between WT and *Nfya*-cKO. Similarly, we collected secondary follicle oocytes at postnatal day 10.

Female mice (7-8 weeks) were super-ovulated through an initial injection of 5 IU pregnant mare serum gonadotropin (PMSG, BioVendor, RP1782725000), followed by a 5 IU injection of human chorionic gonadotropin (hCG, Sigma, C1063) 48 hours later. Oocyte-cumulus complexes were collected 14 hours post hCG injection. Sperms were collected from the cauda epididymis of adult male mice (8-12 weeks). The sperm suspension was capacitated for 1 hour in 200 μl HTF medium (Millipore, MR-070-D). Oocytes were then incubated with spermatozoa for 6-hours. The time when sperms were added to oocytes was considered as 0 hpf. Two-nuclear zygotes were cultured in the KSOM medium (Millipore, MR-106-D) under a humidified atmosphere of 5% CO_2_ at 37°C for further development.

### dTAG13 treatment

dTAG13 (Tocris, 6605) was reconstituted in DMSO to a 50 mM stock. For *Nfya*^dTAG/dTAG^ mESCs treatment, dTAG13 was dilute in mESCs culture medium to 0.5 μM. For *Nfya*^dTAG/dTAG^ embryos treatment, dTAG13 was dilute in KSOM to 0.5 μM. Embryos were washed with KSOM with dTAG13 for at least three times, then cultured in KSOM with dTAG13 for further development.

### Chaperone inhibitor treatment

Ganetespib (MCE, HY-15205) and VER155008 (Selleckchem, S7751) were reconstituted in DMSO to a 100 mM or 150 mM stocks, respectively. For embryos treatment, Ganetespib and VER155008 were dilute in KSOM to 5 nM and 5 μM, respectively. Embryos were washed with KSOM with inhibitors for at least three times, then cultured in KSOM with Ganetespib and VER155008 for further development. L2C embryos were collected for RNA-seq. For ovary treatment, P4 ovaries were dissected and cultured in media [DMEM-F12 media supplemented with 3 mg/ml bovine serum albumin (BSA), 10% fetal bovine serum (FBS) and 10 mIU/ml follicular stimulating hormone (FSH)] with or without 5 nM Ganetespib and 5 μM VER155008 for further development. Growing oocytes (diameter > 25 μm) from 6 days cultured P4 ovaries were collected for RNA-seq.

### Western blot

Cells were lysed in 1 x LDS buffer (Invitrogen, NP0007) at 1 × 10^4^ cell/μL final concentration and heated at 100 °C for 10 min. Samples were run on NuPAGE 4-12% gel (Invitrogen, NP0322BOX) and transferred onto PVDF Transfer Membrane. Primary antibodies used included anti-NFYA (1:250, sc-17753X, lot#D0522, Santa Cruz Biotechnology), anti-HA (1:1000, 3724S, Cell Signaling Technology), and anti-GAPDH (1:50000, 60004-1-Ig, Proteintech). Protein bands were detected with ECL kit (Thermo Fisher Scientific, 32209) and imaged by iBright 1500.

### FerroOrange and C11 staining

Staining procedures were based on the manufacturer’s manual. Briefly, oocytes were incubated in M2 medium (Sigma, M7167) with 1 mM FerroOrange (Dojindo, F374) and 1 μg/mL Hoechst 33342 (Invitrogen, H3570) for 30 min at 37°C. For imaging, oocytes were maintained in the M2 medium with 1 mM FerroOrange. For C11 staining, oocytes were incubated in M2 medium (Sigma, M7167) with 5 μM BODIPY™ 581/591 C11 (Invitrogen, D3861) and 1 μg/mL Hoechst 33342 (Invitrogen, H3570) for 30 min at 37°C before imaged in M2 medium.

### ER staining

The procedures were based on the manufacturer’s manual. Briefly, oocytes were incubated in M2 medium (Sigma, M7167) with ER-ID (1:1000, Enzo Life Sciences, ENZ-51025) and 1 μg/mL Hoechst 33342 (Invitrogen, H3570) for 30 min at 37°C before imaged in M2 medium.

### *In vitro* follicle culture

The experiments were performed as previously described with minor modifications ^94^. Briefly, ovaries were removed from 10-day-old female mice and transferred to dissection medium (Leibovitz L-15 (Gibco, 11415064). Follicles were mechanically isolated with 30G needles in dissection medium supplemented with 1% FBS (Hyclone, SH3007003) and 0.5% penicillin-streptomycin (Gibco, 15140122)). Isolated follicles were washed and recovered in maintenance medium [MEMα with GlutaMAX (Gibco, 32561037) supplemented with 1% FBS (Hyclone, SH3007003) and 0.5% penicillin-streptomycin (Gibco, 15140122)] for 1 h at 37°C under 5% CO. High-quality follicles were selected and encapsulated in 0.5% alginate hydrogels via a droplet-based method. Briefly, follicles were suspended in alginate (Sigma, 71238) solution and droplets (∼5 μL) were crosslinked in 50 mM CaCl (Sigma, 21115) for 2 min. Encapsulated follicles were briefly equilibrated and then cultured in growth medium [α-MEM/F12 (Gibco, 31765035 and 32561037) supplemented with 3 mg/mL bovine serum albumin (Jackson ImmunoResearch, 001-000-162), 1% insulin-transferrin-selenium (Gibco, 41400045), 10 mIU/mL recombinant follicle-stimulating hormone (Sigma, F4021), and 5 mg/mL fetuin (Sigma, F3385)] at 37°C under 5% CO. Half of the medium was changed every 48 hours with 10 μM Ferrostatin-1 (Sigma, SML0583) or an equivalent volume of DMSO (Sigma, 472301) added to the medium.

### Immunofluorescence

Oocytes, embryos, or ovaries were fixed in 4% paraformaldehyde at room temperature 20min or 4 overnight and permeabilized in 0.5% Triton X-100-0.1% Polyvinylalcohol - PBS (PBST-PVA) at room temperature for 1 h. After blocking with 1% BSA in PBST-PVA, the samples were incubated in primary antibodies diluted in PBST-PVA containing 1% BSA overnight at 4. Antibodies were used as follows: anti-NFYA (1:200, sc-17753X, lot#D0522, Santa Cruz Biotechnology), anti-HA (1:200, 3724S or 2367S, Cell Signaling Technology), anti-caspase3 (1:500, 9664S, Cell Signaling Technology), anti-γH2AX (1:200, 05-636-25UG, Millipore), anti-LC3 (1:1000, 4108S, Cell Signaling Technology), anti-DDX4 (1:200, 8761T, Cell Signaling Technology). Following three washes with PBST-PVA, the samples were incubated with appropriate Alexa 488 or 594 fluorophores conjugated secondary antibodies for 1 h at room temperature. Finally, the samples were counterstained with DAPI. The fluorescence signals were imaged using a confocal laser scanning microscope (Zeiss, LSM800). The fluorescent intensity was quantified using ImageJ. For oocyte quantification in the ovaries after *in vitro* culture, growing oocytes were defined by counting DDX4 positive oocytes bigger than 25 μm in diameter that were surrounded by cuboidal granulosa cells. Growing oocytes in the whole ovary were counted by imageJ analysis software.

### Hematoxylin and eosin (HE) staining and electron microscopy

The ovary samples were fixed with 4% paraformaldehyde overnight. The fixed placentas were sent to the Rodent Histopathology Core (220 Longwood Ave., Boston, MA, USA) for paraffin embedding, section and HE staining. Electron microscopic imaging was done by the HMS Electron Microscopy Core.

### CUT&RUN, ATAC and RNA-seq library preparation and sequencing

CUT&RUN assays were performed as previously described with minor modifications ^31^. For mESCs CUT&RUN with more than 1 × 10^4^ cells, cells were resuspended in 50 μl washing buffer (20 mM HEPES/pH=7.5, 150 mM NaCl, 0.5 mM spermidine and 1× protease inhibitor) with activated Concanavalin A Magnetic Beads (Polysciences, 86057-3) for 10 mins at room temperature (RT), then samples were incubated with anti-NFYA (1:100, Invitrogen, MA5-36198, lot#2L4586018) overnight at 4°C. Samples were then incubated with 2.8 ng/μl pA-MNase (home-made) for 2 hrs at 4 °C. Subsequently, samples were incubated with 200 μl pre-cooled 0.5 μM CaCl2 for 20 mins at 4 °C and quench by the addition of 23 μl 10×stop buffer (1.7 M NaCl, 20 mM EGTA, 100 mM EDTA, 0.02% Digitonin, 250 µg/ml glycogen and 250 µg/ml RNase A). DNA fragments were released by incubation at 37°C for 15 mins. Then, 2.5 μl 10% SDS and 2.5 μl 20 mg/ml Protease K (Thermo Fisher) was added and incubated at 55 °C for at least 1 h. DNA was extracted by phenol-chloroform followed by ethanol precipitation. For low input mESCs and embryo CUT&RUN, some modifications were made. Briefly, mESCs, zona-free embryos or isolated oocytes were resuspended in 50 μl wash buffer with activated Concanavalin A Magnetic Beads for 10 mins at RT, then samples were incubated with anti-NFYA overnight at 4°C. The subsequent procedures are the same as those for mESCs described above. Sequencing libraries were prepared with the NEBNext Ultra II DNA library preparation kit for Illumina (New England Biolabs, E7645S).

ATAC-seq was performed as previously described with some modifications ^22,31,35^. Briefly, isolated oocytes were digested with adapter-loaded Tn5 for 15 mins at 37°C, and stopped by stop buffer (100 mM Tris/pH=8.0, 100 mM NaCl, 40 µg/ml Proteinase K and 0.4% SDS) and incubated overnight at 55°C. Then, 5 µl of 25% Tween-20 was added to quench SDS. Sequencing libraries were prepared with NEBNext High-Fidelity 2×PCR Master Mix (NEB, M0541S).

RNA-seq library was prepared by SMART-Seq® Stranded Kit (Takara Bio, 63444) using 5,000 mESCs, 20∼50 isolated oocytes, and ∼10 L2C embryos. All libraries were sequenced by NextSeq 550 system (Illumina) with paired-ended 75-bp reads (**Table S9**)

### RNA-seq data processing

Raw sequencing reads were trimmed using Trimmomatic ^95^ (v0.39) to remove sequencing adaptors, and subsequently mapped to the GRCm38 genome using STAR ^96^ (v2.7.8a). Gene expression was quantified with featureCounts ^97^ (v2.0.1) by counting reads mapped to each gene. Then, edgeR ^98^ (v3.32.1) was employed for normalization and differential expression analysis. Genes with RPKM lower than 1 were defined as lowly expressed genes and excluded from the differential expression analysis. Differentially expressed genes (DEGs) were identified using likelihood ratio test (glmFit and glmLRT functions from edgeR). Genes with a false discovery rate (FDR) below 0.05 and an absolute value of fold change greater than 2 were defined as DEGs. Genes upregulated during primordial to primary follicle transition were classified as “early-PFA genes”, while those upregulated from primary to secondary follicle transition were designated as “late-PFA genes” (**Table S2**). The minor and major ZGA genes were defined as previously described ^31,99^ (**Table S2**). Gene Ontology (GO) enrichment was performed using the R package clusterProfiler ^100^. Gene Set Enrichment Analysis (GSEA) was performed using clusterProfiler and enrichplot. The gene set of Ferroptosis Suppressors was obtained from Ferrdb (http://www.zhounan.org/ferrdb/current/).

### CUT&RUN data analysis

Raw sequencing reads were trimmed using Trimmomatic ^95^ (v0.39) to remove sequencing adaptors, and subsequently aligned to the GRCm38 reference genome using bowtie2 ^101^ (v2.4.2) with parameters: --local --very-sensitive-local --no-unal --no-mixed --no-discordant --dovetail -I 10 -X 700 --soft-clipped-unmapped-tlen. PCR duplicates were removed by Picard MarkDuplicates ^102^ (v2.23.4) and reads with a mapping quality below 30 were removed. The mapped reads were further filtered to only retain proper paired reads with fragment length between 10 and 120 bp. Then, MACS2 ^103^ (v2.2.7.1) was used to call significant peaks with parameters “-f BAMPE -B --SPMR -q 0.05 -g mm --keep-dup all”. To obtain highly conserved NFYA binding sites, only peaks present in both biological replicates were defined as binding sites. The signal tracks were generated with deeptools ^104^ bamCoverage (v3.5.1) with bin size of 1 and normalized using CPM. Peak annotation was performed using the ChIPseeker ^105^ R package. Peaks located within ±2,500 bp of transcription start sites (TSS) were classified as promoter peaks, whereas peaks outside this region were considered distal peaks. The heatmaps of binding profiles were calculated with deeptools ^104^ computeMatrix (v3.5.1) using bigwig signal tracks as input and bin size of 10 and visualized in R with profileplyr and EnrichedHeatmap ^106^ packages.

### ATAC–seq data analysis

ATAC–seq data were processed with the ENCODE ATAC–seq pipeline using default parameters (v2.1.2, https://github.com/ENCODE-DCC/atac-seq-pipeline). Peaks located within ±2,500 bp of transcription start sites (TSS) were classified as promoter peaks, whereas peaks outside this region were considered distal peaks.

To identify differentially accessible peaks, the ATAC-seq read counts within each NFYA-binding site were quantified using multiBamCov ^107^ (v2.30.0). The resulting read count matrix was imported into edgeR ^98^ for downstream analysis. After normalization and dispersion estimation, a generalized linear model was fitted, and likelihood ratio tests were performed (glmFit and glmLRT functions). Peaks with a false discovery rate (FDR) less than 0.05 were defined as differentially accessible peaks.

### Motif enrichment analysis

Motif enrichment analysis was performed using HOMER ^108^ (v4.11) findMotifsGenome.pl with the mm10 reference and parameter -size 200, using peak regions as input. Briefly, HOMER identifies over-represented motifs in the input peak regions by comparing them to matched random background sequences. Enrichment P values for all motifs in all peaks were then corrected for multiple testing by calculating adjusted P value using the Benjamini-Hochberg method. Motifs with an adjusted p-value less than 0.05 were considered significantly enriched. The full spectrum of highly enriched TF motifs identified in Figure 1D and S1D are included in **Table S10**. The enriched TF motifs identified in Figure 6B are also included in **Table S10.**

### Motif occurrence analysis

Motif occurrence analysis was performed using HOMER ^108^ (v4.11) annotatePeaks.pl with parameters mm10 -size −500,500 -hist 20 -ghist for regions around gene TSSs. Motif files were downloaded from the JASPAR database14 and manually converted to HOMER motif format. The JASPAR ^109^ motif ID for TFs analyzed in this study is NFYA (MA0060.1) and OBOX (PH0121.1), and a log odds detection threshold of 6.0 was applied. The motif occurrence matrix was visualized in R with the EnrichedHeatmap ^106^ package.

### Prediction of NFYA-mediated promoter-enhancer interactions

Multiple public dataset in the 2-cell stage were collected as the input for Activity-by-Contact (ABC) model (v0.2) ^66^, including, RNA-seq, ATAC-seq ^33^, H3K27ac CUT&RUN ^99^, and L2C NFYA CUT&RUN, to predict NFYA-mediated promoter–enhancer interactions. Enhancers within 5 Mb and with scores ≥0.02 were assigned to target genes.

### Quantification and Statistical Analysis

Statistical analyses were performed in GraphPad Prism 10 (v10.2.0) or R (v4.3.2) with the exceptions of GSEA data, where analysis was performed using the GSEA software (v4.3.2) and GO term analysis where analysis was performed using the clusterProfiler. Correlation was calculated using Pearson’s correlation coefficient. Unless otherwise stated, comparisons between two groups were made using a two-tailed, unpaired Student’s *t*-test. Where multiple comparisons were made, these were corrected using Benjamini-Hochberg False Discovery Rate (FDR) adjustment. *n* refers to the number of independent biological replicates. *P*-values are denoted as ^∗^*P* < 0.05 and ^∗∗^*P* < 0.01, between the indicated groups. “ns” = “not significant”. Changes were considered to be statistically significant when *P* (or FDR where appropriate) < 0.05. Where biological replicate data are presented in plots, these are given as mean ± standard error of the mean.

## Supplemental tables (separated files)

**Table S1.** Differential expressed genes in primary follicle oocytes of *Nfya*-cKO versus WT, Related to Figure 2.

**Table S2.** Gene lists used in this study, Related to STAR Methods.

**Table S3.** Differential expressed genes in secondary follicle oocytes of *Nfya*-cKO versus WT, Related to Figure 3.

**Table S4.** Differentially expressed genes of late two-cell embryos of NFYA^dTAG/dTAG^ treated with dTAG13 versus DMSO, Related to Figure 5.

**Table S5.** Differentially expressed genes of late two-cell embryos of WT treated with G&V versus DMSO, Related to Figure 7.

**Table S6.** Differential expressed genes of growing oocytes from 6 days in vitro cultured P4 ovaries of WT treated with G&V versus DMSO, Related to Figure 7.

**Table S7.** Oligos used in this study, Related to STAR Methods.

**Table S8.** ddPCR validated Nfya-dTAG knock-in copy number, Related to STAR Methods.

**Table S9.** Summary of the sequenced libraries in this study, Related to STAR Methods.

**Table S10.** List of the full spectrum of highly enriched TF motifs identified in Fig. 1D, S1D, and 6B, Related to STAR Methods.

