## Supplementary material for "NFYA regulates two genome-wide transcriptional activation events during oocyte-to-embryo transition": Figure. S1 to 7

### Supplemental Figure titles and legends

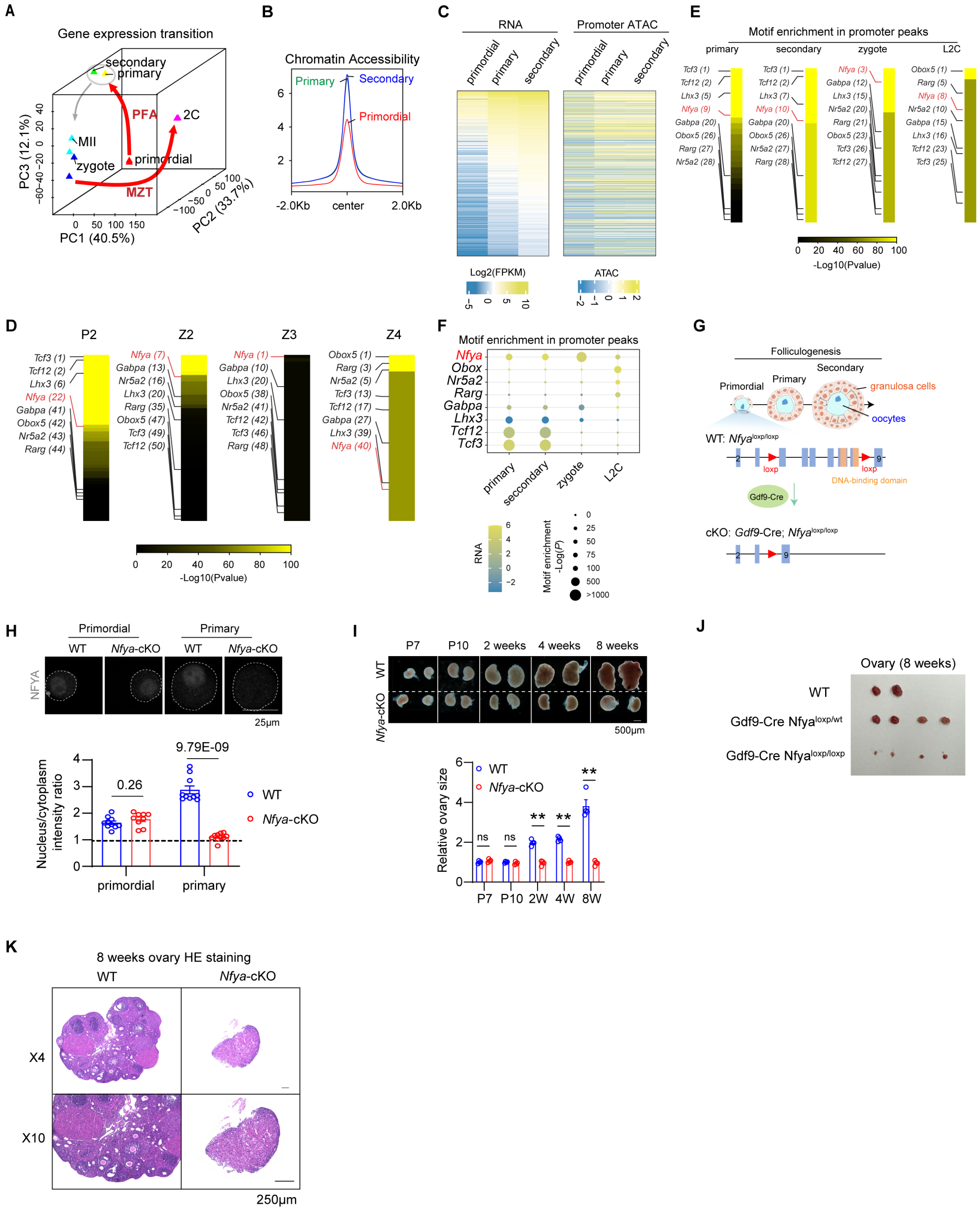


**Figure S1. Identification of NFYA as a potential regulator of PFA and ZGA and demonstration that NFYA loss impairs folliculogenesis, Related to Figure 1. (A)** Principal components analysis (PCA) of gene expression from primordial follicle to late 2-cell embryos.

**(B)** Metaplot showing the fold change of ATAC–seq signals (ATAC_FC) around transcription start sites (TSSs) of primordial, primary, and secondary follicles. **(C)** Heatmaps showing the expression of upregulated genes from primordial to primary and secondary follicles, and the corresponding ATAC_FC signals at their promoters. **(D)** Enrichment of top TF motifs at the ATAC–seq peak groups (P2, Z2, Z3, and Z4) during PFA and MZT. **(E-F)** Enrichment of top TF motifs (E) and representative TF (F) at promoter ATAC–seq peaks of the indicated stages. **(G)** Schematic diagrams showing the early folliculogenesis and a conditional *Nfya* knockout strategy using *Gdf9*-Cre-mediated deletion of exons 4-8 harboring its DNA-binding domain. *Gdf9*-Cre is specifically active in oocytes starting from primordial follicle stage. **(H)** Upper panel: the grey scale of NFYA signals in WT and Gdf9-Cre *Nfya*$^{loxp/loxp}$ (*Nfya*-cKO) oocytes from figure 1G. Scale bar, 25 μm. Lower panel: relative nuclear to cytoplasmic intensity ratio of NFYA in primordial and primary follicle oocytes from WT and *Nfya*-cKO female mouse. P values, Student's t test. **(I)** Upper panel: The gross morphology of ovaries derived from WT and *Nfya*-cKO mice at postnatal day 7 (P7), 10, 2 weeks, 4 weeks, and 8 weeks. Lower panel: the relative size of ovaries from WT (n=4) and *Nfya*-cKO mice (n=4). Scale bar, 500 μm. **(J)** Images of 8-week ovaries from WT (*Nfya*$^{loxp/loxp}$), Gdf9-Cre *Nfya*$^{loxp/wt}$, and Gdf9-Cre *Nfya*$^{loxp/loxp}$ (*Nfya*-cKO) mice. **(K)** H&E staining of paraffin embedded 8-week ovarian sections of WT and *Nfya*-cKO mice. Scale bar, 250 μm.

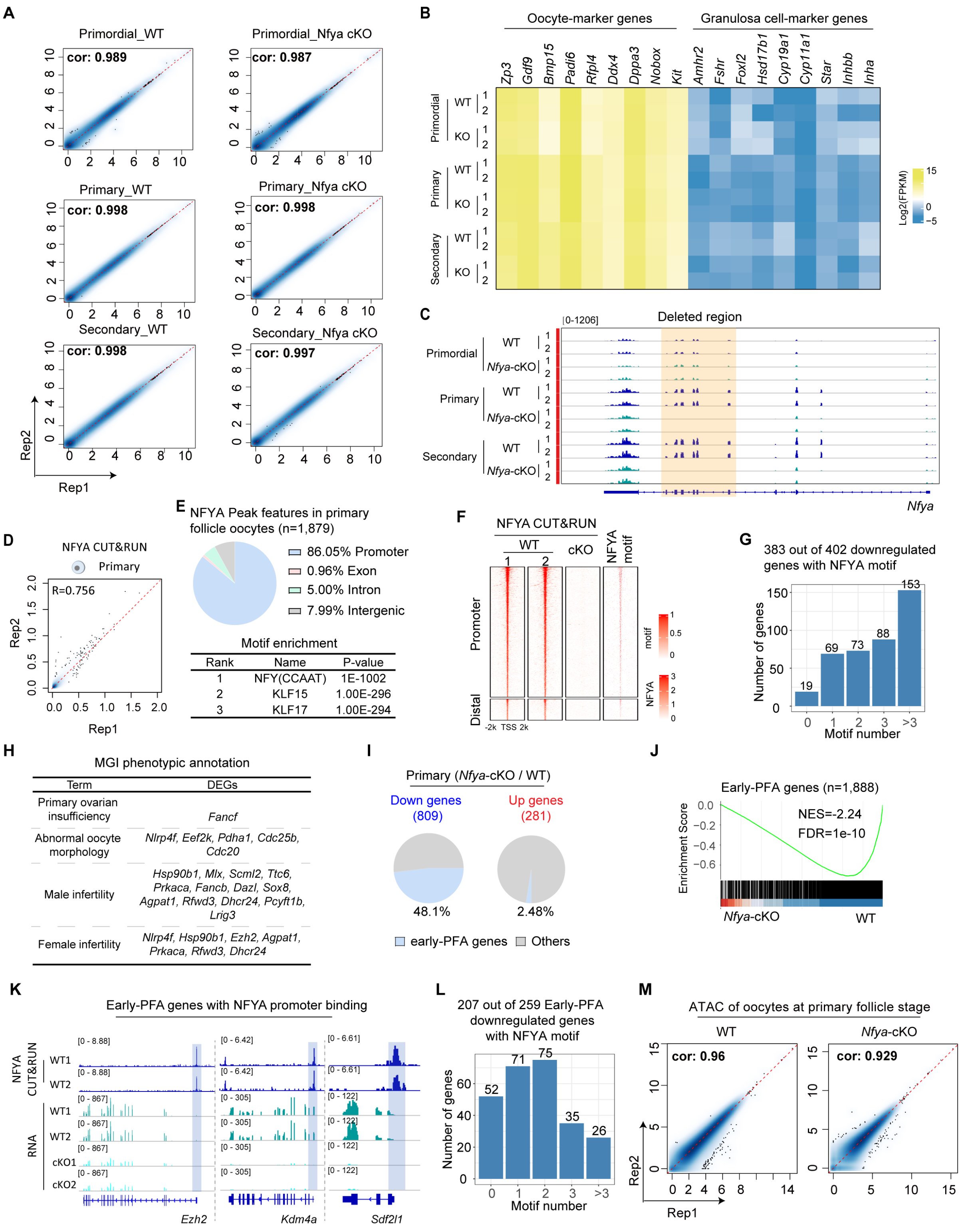


**Figure S2. NFYA deficiency impairs PFA in primary follicle oocytes, Related to Figure 2.** **(A)** Correlations of the replicates of the total RNA-seq. The x and y axis of the dot plots are Log (CPM+1). **(B)** Heatmap showing the expression of oocyte- and granulosa cell-marker genes in the WT and *Nfya*-cKO primordial, primary, and secondary follicle oocytes. **(C)** Genome browser view of RNA-seq at the *Nfya* genomic regions in WT and *Nfya*-cKO oocytes at primordial, primary, and secondary follicle stages. **(D)** Correlation of replicates for NFYA CUT&RUN in primary follicle oocytes. Each dot represents a 5-kb bin. **(E)** Upper panel: the genomic distribution of NFYA binding peaks generated by NFYA CUT&RUN in primary follicle oocytes. Lower panel: the top 3 TF motifs enriched in the regions bound by NFYA in primary follicle oocytes. Note that based on the p-value, the NFYA motif is much more highly enriched compared to the other enriched motifs. **(F)** Heatmaps showing all the CUT&RUN signals of NFYA peaks at promoters and distal regions in WT or *Nfya*-cKO primary follicle oocytes. **(G)** NFYA motif numbers found in the promoters of the other 402 downregulated genes in primary follicle oocyte. **(H)** MGI phenotypic annotation for NFYA-bound downregulated genes (n=402) in primary follicles. Infertility related terms and example genes are shown. **(I)** The percentage of differentially expressed (*Nfya*-cKO vs WT) early-PFA genes in primary follicle oocytes. **(J)** Gene set enrichment analysis (GSEA) showing the early-PFA genes (n=1,888) are significantly decreased in *Nfya*-cKO oocytes. **(K)** Genome browser view of NFYA targeted early-PFA gene examples of NFYA CUT&RUN and RNA-seq signals in primary follicles. **(L)** NFYA motif numbers found in the promoters of the other 259 downregulated genes in early-PFA. **(M)** Correlation of replicates for ATAC-seq in WT or *Nfya*-cKO primary follicle oocytes. The x and y axis of the dot plots are log2(normalized_counts+1). Each dot represents a 5-kb bin.

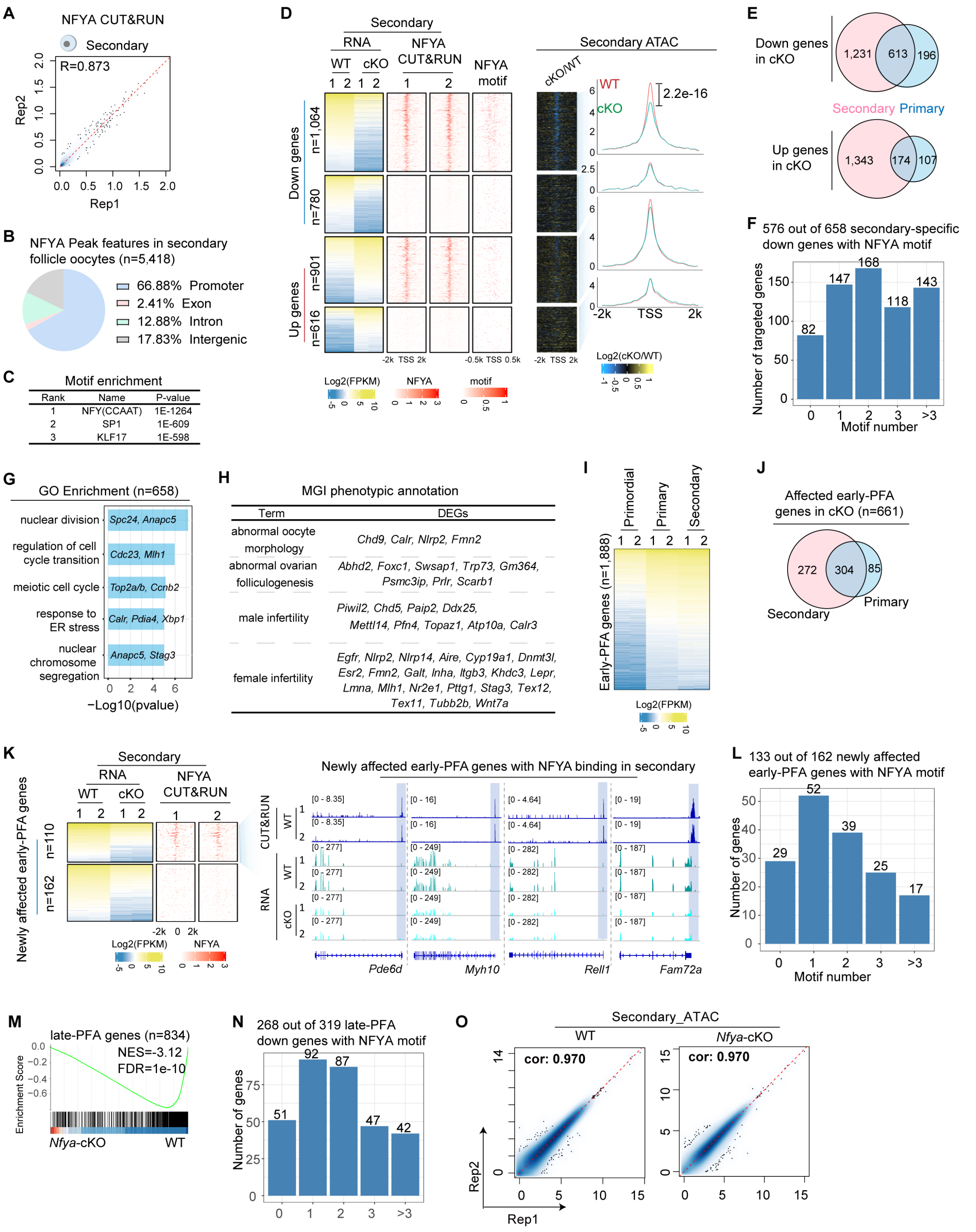


**Figure S3. NFYA is critical for PFA in secondary follicle oocytes, Related to Figure 3. (A)** Correlation of replicates for NFYA CUT&RUN in secondary follicle oocytes. The x and y axis of the dot plots are log2(normalized_counts+1). Each dot represents a 5-kb bin. **(B)** The genomic distribution of NFYA binding peaks generated by NFYA CUT&RUN in secondary follicle oocytes. **(C)** The top 3 TF motifs enriched in the regions bound by NFYA in secondary follicle oocytes. **(D)** Left panel: heatmaps showing the gene expression, NFYA CUT&RUN signals, NFYA motif occurrence, and ATAC–seq signal fold change (ATAC_FC) in the promoter regions of the DEGs in the secondary follicle oocytes upon NFYA loss. The downregulated or upregulated genes are separated into two groups based on whether they have direct NFYA promoter binding. Right panel: density plots showing the average ATAC–seq signal fold change (cKO/WT) in the promoter regions of the DEGs in different groups of the secondary follicle oocytes. P values are calculated with two-sided Student's t-test. **(E)** A comparison of the downregulated and upregulated genes between primary follicle oocytes and secondary follicle oocytes after *Nfya* knockout. **(F)** NFYA motif numbers found in the promoters of the other 658 downregulated genes in secondary follicle oocytes. **(G)** GO terms enriched in the NFYA-bound down-regulated genes (n=658) in the secondary follicle oocytes. The example genes are shown in each GO term. **(H)** MGI phenotypic annotation for NFYA-bound newly downregulated genes (n=658) in secondary follicle oocytes. Infertility related terms and gene examples are shown. **(I)** Heatmaps showing the gene expression of early-PFA genes (n=1,888) in oocytes at primordial, primary, and secondary follicle stages. **(J)** A comparison of the downregulated early-PFA genes between primary follicle oocytes and secondary follicle oocytes after *Nfya* knockout. **(K)** Left panel: heatmaps showing the newly downregulated early-PFA genes (n=272) in secondary follicle oocytes upon NFYA loss, and the NFYA binding around the TSS of the corresponding genes. The downregulated genes are separated into two groups based on whether they have direct NFYA promoter binding. Right panel: Genome browser view of NFYA targeted newly affected early-PFA gene examples of NFYA CUT & RUN and RNA-seq in secondary follicle oocytes. **(L)** NFYA motif numbers found in the promoters of the 162 newly affected early-PFA genes. **(M)** GSEA enrichment plots showing the late-PFA genes (n=834) are significantly decreased in the *Nfya*-cKO oocytes. **(N)** NFYA motif numbers found in the promoters of the 319 downregulated late-PFA genes. **(O)** Correlation of replicates for ATAC-seq in WT and *Nfya*-cKO secondary follicle oocytes. The x and y axis of the dot plots are log2(normalized_counts+1). Each dot represents a 5-kb bin.

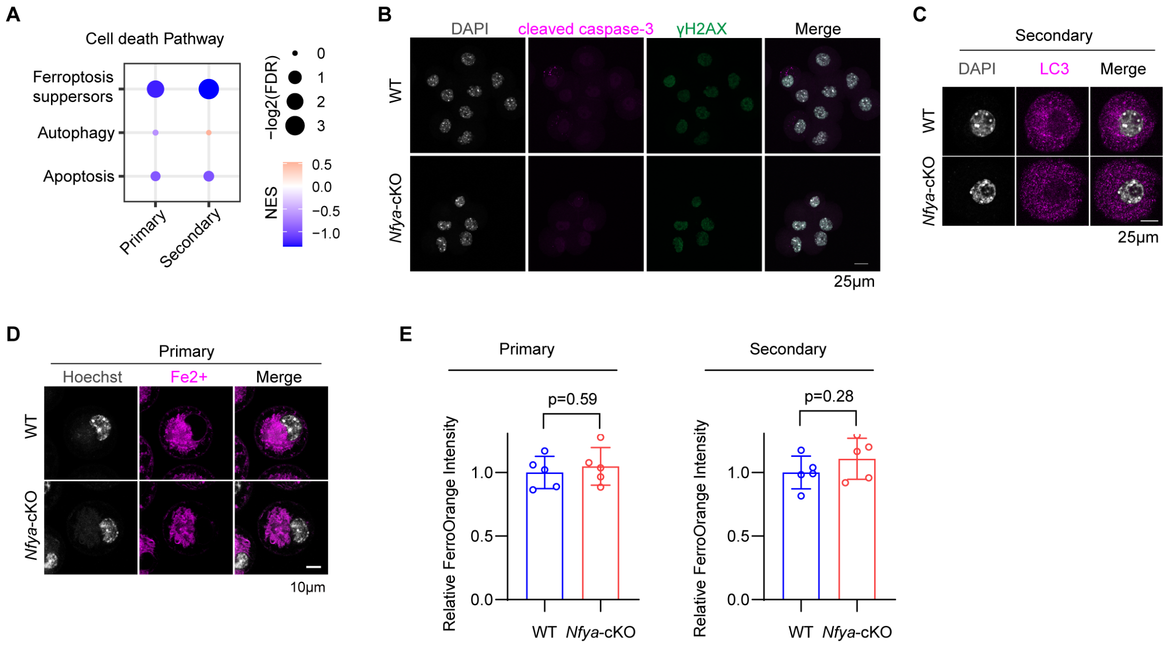


**Figure S4. NFYA loss triggers non-canonical ferroptosis in growing oocytes, Related to Figure 4. (A)** Dotplots of GSEA enrichment showing pathways of ferroptosis, but not apoptosis or autophagy, are enriched in *Nfya*-cKO oocytes. **(B)** Cleaved caspase3 (red) and γH2AX (green) staining show no sign of apoptosis in *Nfya*-cKO oocytes. Scale bar, 25 µm. **(C)** LC3 (red) staining does not show sign of autophagy in *Nfya*-cKO secondary follicle oocytes. Scale bar, 25 µm. **(D)** Representative images of FerroOrange (an $Fe^{2+}$ indicator) staining of WT and *Nfya*-cKO primary follicle oocytes. Scale bar, 10 µm. **(E)** Relative intensity of FerroOrange staining in primary and secondary follicle oocytes. P values, Student's t test.

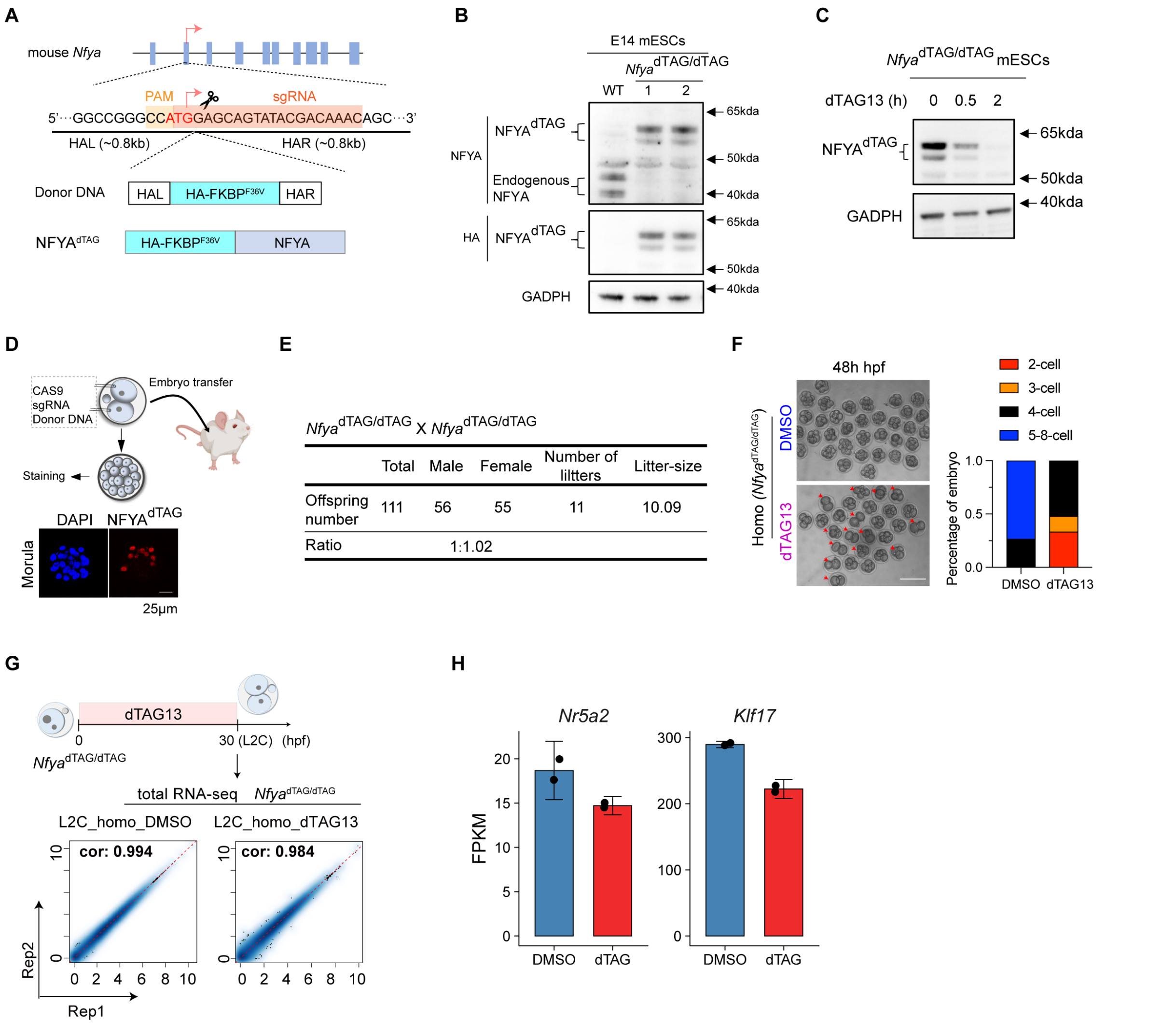


**Figure S5. Generation of NFYA$^{dTAG}$ mice and NFYA depletion affects major ZGA, Related to Figure 5. (A)** Diagram illustration of HA-FKBP (F36V) knock-in at the 5' end of *Nfya* gene and the NFYA$^{dTAG}$ fusion protein. **(B)** Western blot analysis confirming NFYA$^{dTAG}$ knock-in in ES cells. The blots were incubated with anti-NFYA and anti-HA, respectively. GAPDH was used as a loading control. **(C)** Western blot analysis confirming dTAG13-mediated NFYA degradation in mES cells. The blots were incubated with anti-HA. GAPDH was used as a loading control. **(D)** Diagram illustration for *Nfya*$^{dTAG/dTAG}$ mouse generation and HA staining confirming the dTAG knock-in cells in morula. Scale bar, 25 μm. **(E)** Statistics of pup numbers from NFYA$^{dTAG/dTAG}$ × NFYA$^{dTAG/dTAG}$ crosses, indicating normal function of the NFYA$^{dTAG}$ fusion protein. **(F)** Representative images (right panel) and developmental rates (left panel) of 48 hpf (4-8 cell stage) Homo embryos with or without dTAG13 treatment. **(G)** Upper panel: schematic diagram of *Nfya*$^{dTAG/dTAG}$ embryo generation and dTAG13 treatment. Lower panel:

correlation of replicates for total RNA-seq in *Nfya*$^{dTAG/dTAG}$ L2C embryos with DMSO or dTAG13 treatment. The x and y axis of the dot plots are Log (CPM+1). **(H)** *Nr5a2* and *Klf17* expression in *Nfya*$^{dTAG/dTAG}$ L2C embryos with or without dTAG13 treatment.

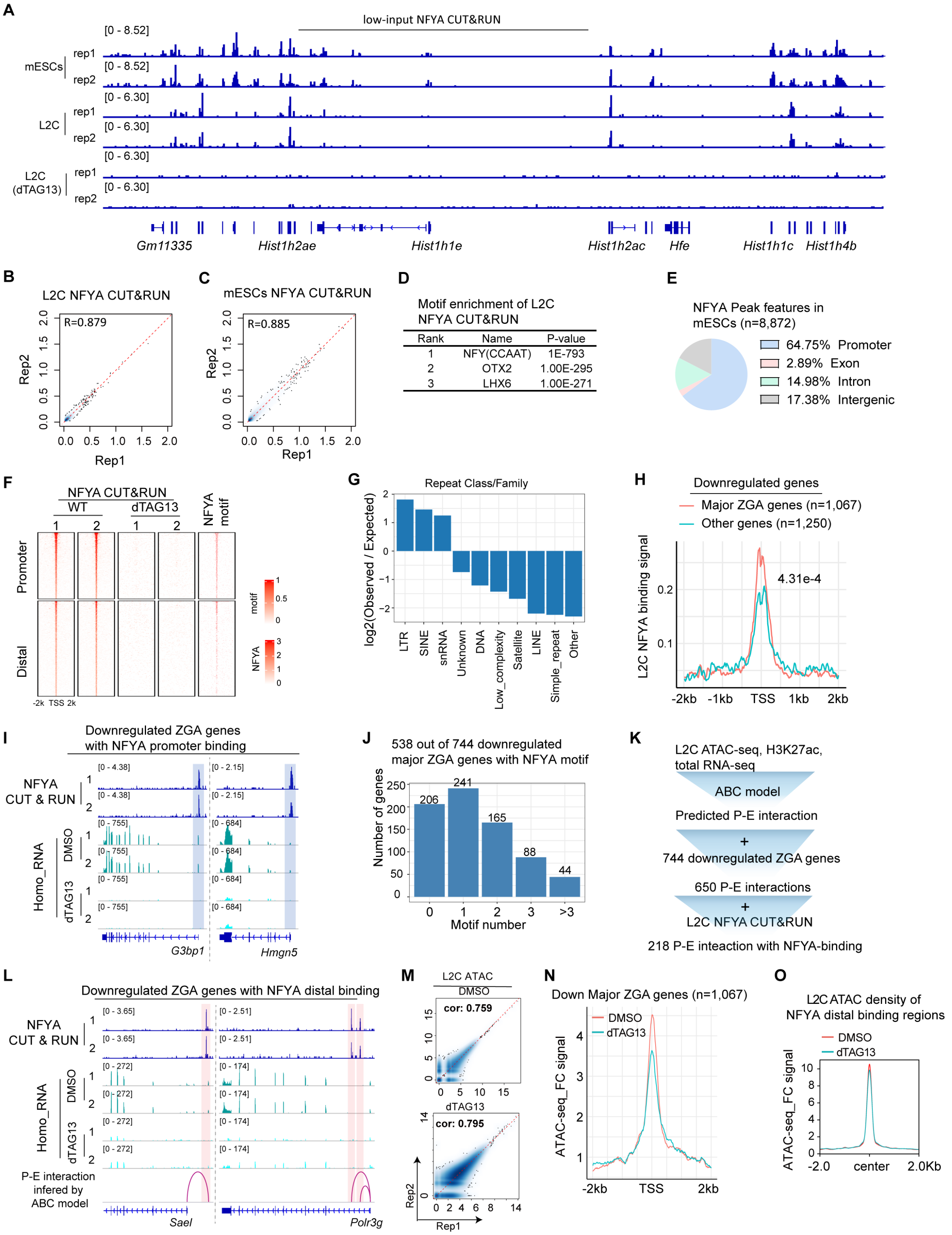


**Figure S6. NFYA affects ZGA by promoter and distal binding, Related to Figure 5. (A)** Genome browser view showing examples of NFYA CUT&RUN profiles in mESCs (1000 cells), WT L2C embryos, and maternal NFYA-depleted L2C embryos by dTAG13 (L2C-dTAG13). **(B)**

Correlation between replicates of NFYA CUT&RUN in late 2cell embryos. The x and y axis of the dot plots are log2(normalized_counts+1). Each dot represents a 5-kb bin. **(C)** Correlation between replicates of NFYA CUT&RUN in mESCs. The x and y axis of the dot plots are Log2(normalized_counts+1). Each dot represents a 5-kb bin. **(D)** The top 3 TF motifs enriched in the regions bound by NFYA in L2C embryos. **(E)** The genomic distribution of NFYA binding peaks from NFYA CUT&RUN in mESCs. **(F)** Heatmaps showing all the CUT&RUN signals of NFYA peaks at promoters and distal regions in L2C with or without dTAG13 treatment. **(G)** A bar chart showing repeat class/family enrichment at NFYA-binding peaks in the distal regions of L2C embryos. **(H)** Density plots showing the average L2C NFYA CUT&RUN signal in the promoter regions of downregulated major ZGA genes (n=1,067) and non-ZGA genes (n=1,250) at late 2-cell stage upon NFYA depletion. P values were calculated with two-sided Student's t-test. **(I)** Genome browser view of NFYA promoter targeted ZGA gene examples of NFYA CUT & RUN and RNA-seq in late 2-cell embryos. **(J)** NFYA motif numbers found in the promoters of the 744 downregulated ZGA genes. **(K)** Schematic diagram showing prediction of promoter-distal interactions with NFYA distal binding by ABC model. **(L)** Genome browser view of NFYA distal targeted ZGA gene examples of NFYA CUT & RUN and RNA-seq in L2C embryos. **(M)** Correlation of replicates for ATAC-seq in L2C embryos with DMSO or dTAG13 treatment. The x and y axis of the dot plots are log2(normalized_counts+1). Each dot represents a 5-kb bin. **(N)** Density plots showing the average ATAC–seq_FC signal in the promoter regions of the downregulated major ZGA genes of the L2C embryos. **(O)** Density plots showing the average ATAC–seq_FC signal in the NFYA-bound distal regions.

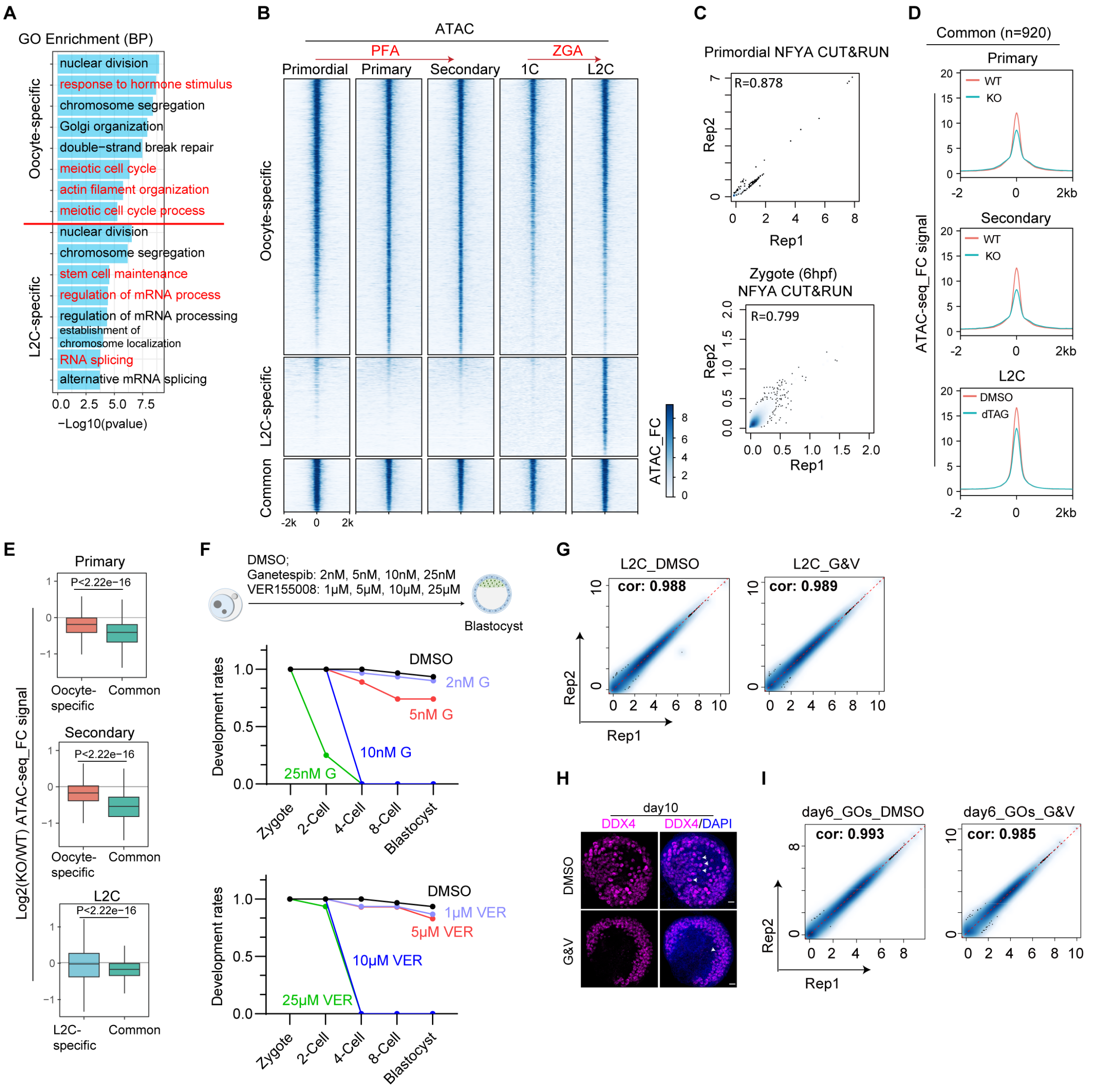


**Figure S7. Conserved and divergent NFYA chromatin bindings regulate PFA and ZGA, Related to Figure 6 and 7. (A)** Representative GO (BP) terms enriched in the oocyte- and L2C-specific bindings related genes. **(B)** Heatmaps showing the dynamics of ATAC–seq signal in the NFYA binding regions during PFA (primordial follicle oocytes, primary follicle oocytes, and secondary follicle oocytes) and ZGA (zygotes and late 2-cell embryo) at promoter and distal regions. **(C)** Correlation between replicates of NFYA CUT&RUN in primordial follicle oocytes and zygotes (6hpf). The x and y axis of the dot plots are log2(normalized_counts+1). Each dot represents a 5-kb bin. **(D)** Density plots showing the average ATAC–seq_FC signals of the 920 NFYA common binding regions in oocytes or L2C embryos after NFYA-depletion. **(E)** Box plots showing the comparison of ATAC–seq_FC signal changes [Log2(KO/WT)] between the oocyte-specific, L2C-specific, and common groups in primary follicle oocytes, secondary follicle oocytes, and L2C embryos. **(F)** Upper panel: schematic diagram of the experimental design. Medium panel: the embryo development rate upon DMSO and Ganetespib: 2nM, 5nM, 10nM, 25nM treatment. Lower panel: the embryo development rate upon DMSO and VER155008: 1μM, 5μM, 10μM, 25μM treatment. **(G)** Correlation of replicates for total RNA-seq in L2C embryos with DMSO or G&V treatment. The x and y axis of the dot plots are Log (CPM+1). **(H)** Ovarian morphology of 10 days *in vitro* cultured P4 ovaries with DMSO or G&V treatment. DDX4 stained oocytes. Scale bar, 50 μm. **(I)** Correlation of replicates for total RNA-seq in growing oocytes (GOs) from P4 ovaries after 6 days DMSO or G&V treatment. The x and y axis of the dot plots are Log (CPM+1).
